# Protection of bona fide T follicular memory cells during tissue isolation reveals their persistence, plasticity and functional impact

**DOI:** 10.1101/677559

**Authors:** Marco Künzli, David Schreiner, Tamara Pereboom, Nivedya Swarnalekha, Jonas Lötscher, Yusuf I. Ertuna, Julien Roux, Florian Geier, Christoph Hess, Justin T. Taylor, Carolyn G. King

**Affiliations:** Immune Cell Biology Laboratory; Department of Biomedicine, University of Basel, University Hospital Basel CH-4031 Basel, Switzerland.; Department of Medicine, CITIID, University of Cambridge, Cambridge, UK; Vaccine and Infectious Disease Division, Fred Hutchinson Cancer Research Center, Seattle, USA; Swiss Institute of Bioinformatics, Basel, Switzerland

## Abstract

CD4 memory T cells play an important role in protective immunity and are a key target in vaccine development. Many studies have focused on T central memory (TCM) cells, while the existence and functional significance of T follicular helper (TFH) memory cells is controversial. Here we show that TFH memory cells are highly susceptible to NAD induced cell death (NICD) during isolation from tissues, leading to their under-representation in prior studies. NICD blockade reveals the persistence of abundant TFH memory cells, with high expression of hallmark TFH markers, that persist to at least 400 days after infection, by which time TCM cells are no longer found. Using single cell RNA-seq we demonstrate that TFH memory cells are transcriptionally distinct from TCM cells, maintain stemness and self-renewal gene expression, and, in contrast to TCM cells, are multipotent following recall. Surprisingly, TFH memory cells concurrently express a distinct glycolytic signature similar to trained immune cells, including elevated expression of mTOR, HIF-1 and cAMP regulated genes. Late disruption of glycolysis/ICOS signaling leads to TFH memory cell depletion concomitant with decreased splenic plasma cells and circulating antibody titers, demonstrating both unique homeostatic regulation of memory TFH and their sustained function during the memory phase of the immune response. These results highlight the metabolic heterogeneity underlying distinct memory T cell subsets and establish TFH memory cells as an attractive target for the induction of long-lived adaptive immunity.

**HIGHLIGHTS:**

- Cell death during isolation from the tissue prevents the full recovery of TFH memory cells
- TFH memory cells are transcriptionally distinct from TCM cells and maintain a broader recall capacity
- TFH memory cells are maintained in the absence of antigen but require ICOS signaling and glycolysis
- TFH memory cells support late phase antibody production by splenic plasma cells

## INTRODUCTION

Vaccination and infection lead to the generation of protective immune responses mediated by both memory T and B cells (Ahmed and Gray, 1996). Two major subsets of memory T cells have been described based on their differential expression of lymphoid homing receptors, including CCR7+ central memory (TCM) cells and CCR7-effector memory (TEM) cells (Sallusto et al., 2004). Following challenge infection, TCM cells produce interleukin-2 and maintain the capacity to proliferate and generate secondary effector cells. In contrast, TEM cells are able to immediately produce inflammatory cytokines but have more limited expansion. In addition to these subsets, CD4+ T follicular helper (TFH) memory cells with the ability to promote secondary B cell expansion and class switching have been described (Asrir et al., 2017; Hale et al., 2013; Luthje et al., 2012; MacLeod et al., 2011; Pepper et al., 2011). Importantly, the number of circulating TFH cells has been shown to correlate with the number of blood plasmablasts following vaccination in humans, and can be boosted to improve long-lived antibody production (Bentebibel et al., 2013; Crotty and Ahmed, 2018; Hill et al., 2019). These data suggest that targeted generation of TFH memory cells may be a rational approach for improving vaccine design. However, despite the importance of TFH cells for supporting productive antibody responses, the signals promoting maintenance and survival of the TFH memory cell compartment are not well understood. In addition, whether or not TFH memory cells retain the capacity to differentiate into diverse secondary effectors is unclear (Hale et al., 2013; Keck et al., 2014; Luthje et al., 2012; Pepper et al., 2011). Analysis of TFH memory is complicated by a gradual loss of phenotypic markers typically associated with TFH effector cells, including PD1 and CXCR5, as well as the apparent decline of the CD4 memory compartment compared to CD8 memory T cells (Hale et al., 2013; Homann et al., 2001; Marshall et al., 2011; Pepper et al., 2011; Williams et al., 2008). Moreover, the relationship between TCM cells and TFH effector cells, which share several surface markers and transcription factors including CXCR5, ICOS, TCF1, STAT3 and ID3, is not well established (Choi et al., 2015; Hale et al., 2013; Ma et al., 2012; Marshall et al., 2015; Siegel et al., 2011; Tian et al., 2017). Recently a TCM precursor signature, including markers for lymphoid homing (Ccr7) and survival (Bcl2), was identified among antigen-specific effector cells responding to viral infection (Ciucci et al., 2019). This signature, however, was not detected in TFH effector cells suggesting an early divergence of TCM and TFH memory precursors. Nevertheless, it remains unclear if TFH cells present at later phases are remnants of a primary effector response, or if they represent a distinct population of self renewing TCM cells that share differentiation requirements and phenotypic characteristics with TFH cells.

In this study we a determined that TFH memory cells are susceptible to NAD induced cell death (NICD) during isolation. By blocking NICD we observed that TFH memory cells are maintained in high numbers to at least 400 days after infection, whereas TCM cells are subject to attrition. Transcriptional and epigenetic profiling revealed that TFH memory cells constitutively engage a glycolytic metabolism while maintaining a stem-like state. Consistent with these findings, transfer experiments revealed the ability of TFH memory cells, but not TCM cells, to generate the full spectrum of secondary effectors. Although TFH memory cells can survive in the absence of antigen, they depend on sustained ICOS signals to preserve glycolytic and Tcf7 dependent gene expression. A reduction in TFH memory cell numbers induced by late ICOS blockade led to a reduction in circulating antibody titers and splenic plasma cells, highlighting an underestimated contribution of TFH memory cells to late phase humoral immune responses.

## RESULTS

### TFH memory cells are susceptible to death during isolation

TFH cells were recently described to express high levels of the purinergic receptor P2X7 receptor (Iyer et al., 2013; Proietti et al., 2014). P2X7R is an ATP-gated cation channel that can be ADP ribosylated by the cell surface enzyme ARTC2.2, rendering certain cell types, including regulatory T cells (Tregs) and resident memory T cells, susceptible to NICD during isolation from the tissue (Aswad et al., 2005; Fernandez-Ruiz et al., 2016). Injection of an ARTC2.2 blocking nanobody (NICD-protector) prior to organ harvest has been shown to protect these subsets from NICD and improve their recovery from lymphoid organs (Borges da Silva et al., 2019; Hubert et al., 2010). To determine whether inhibition of ARTC2.2 could also improve the recovery of TFH cells at effector and memory time points, we harvested antigen-specific T cells from NICD-protector treated mice at various time points after infection with Lymphocytic Choriomeningitis Virus (LCMV). Polyclonal LCMV-specific CD4+ T cells were enriched using tetramer staining for Ia^b^:NP_309-328_ (NP-specific) or IA^b^:GP_66-77_ (GP66-specific) and analyzed for expression of TFH associated surface markers (Hale et al., 2013; Marshall et al., 2011). In untreated mice, TFH effector cells were clearly identified at day 15 after infection, but were largely absent by day 43 (Figures 1A and S1A). In contrast, treatment with NICD-protector resulted in a significant recovery of TFH cells at all time points and with both T cell specificities, indicating a larger expansion and more prolonged survival of TFH cells than previously appreciated (Figures 1A-1C). As the number of GP66-specific TFH cells was approximately 4-fold higher than NP-specific TFH cells, we focused our subsequent analyses on the GP66-specific T cell compartment (Figure S1B). Two-dimensional visualization of the cytometry data by *t-*distributed stochastic neighbor embedding confirmed that NICD-protector preferentially rescued cells with high expression of P2X7R (Figure 1B). NICD-protector also significantly improved the recovery of PSGL1^hi^Ly6C^lo^ (hereafter Ly6C^lo^ Th1) memory cells but had minimal impact on more terminally differentiated PSGL1^hi^Ly6C^hi^ (hereafter Ly6C^hi^ Th1) memory cells, in line with the levels of P2X7R expression on these subsets (Figures 1C, 1E and S1C). In addition, NICD-protector improved the recovery Ly6c^lo^ and CXCR5+ cells after infection with *Listeria monocytogenes*, again in correlation with expression of P2X7R on these subsets (Figures S1E-S1H). After day >400, TFH memory cells were maintained in LCMV infected mice although the mean cell fluorescence of PD1 on this population was decreased compared to earlier time points (Figures 1A and 1C). Unexpectedly, Ly6C^lo^ Th1 memory cells, previously shown to contain a substantial proportion of TCM cells (Marshall et al., 2011), were nearly absent at this time point, suggesting either a survival defect or conversion into one of the remaining memory cell subsets (Figures 1A and S1C). TFH cells isolated at late time points after infection were further phenotyped by flow cytometry and characterized by high expression of FR4, CD73, CXCR4, ICOS and Bcl6 compared to Ly6C^hi^ and Ly6C^lo^ Th1 memory cells (Figures 1D and 1E). Although a similar phenotype was observed on polyclonal NP-specific TFH cells isolated from LCMV infected mice (Figure S1A), monoclonal T cells from SMARTA or NIP T cell receptor (TCR) transgenic strains specific for LCMV GP66:I-A^b^ and NP309:IA^b^ respectively, generated significantly fewer TFH memory cells after LCMV infection (Figure S1D) (Nance et al., 2015; Oxenius et al., 1998). These observations are in agreement with previous reports showing a gradual decline of TFH associated markers on transferred monoclonal populations and highlight the value of studying polyclonal responses, particularly given the tendency of different types of TCR transgenic T cells to undergo distinct and more limited patterns of differentiation (Hale et al., 2013; Marshall et al., 2011; Tubo et al., 2013).

**Figure 1.**
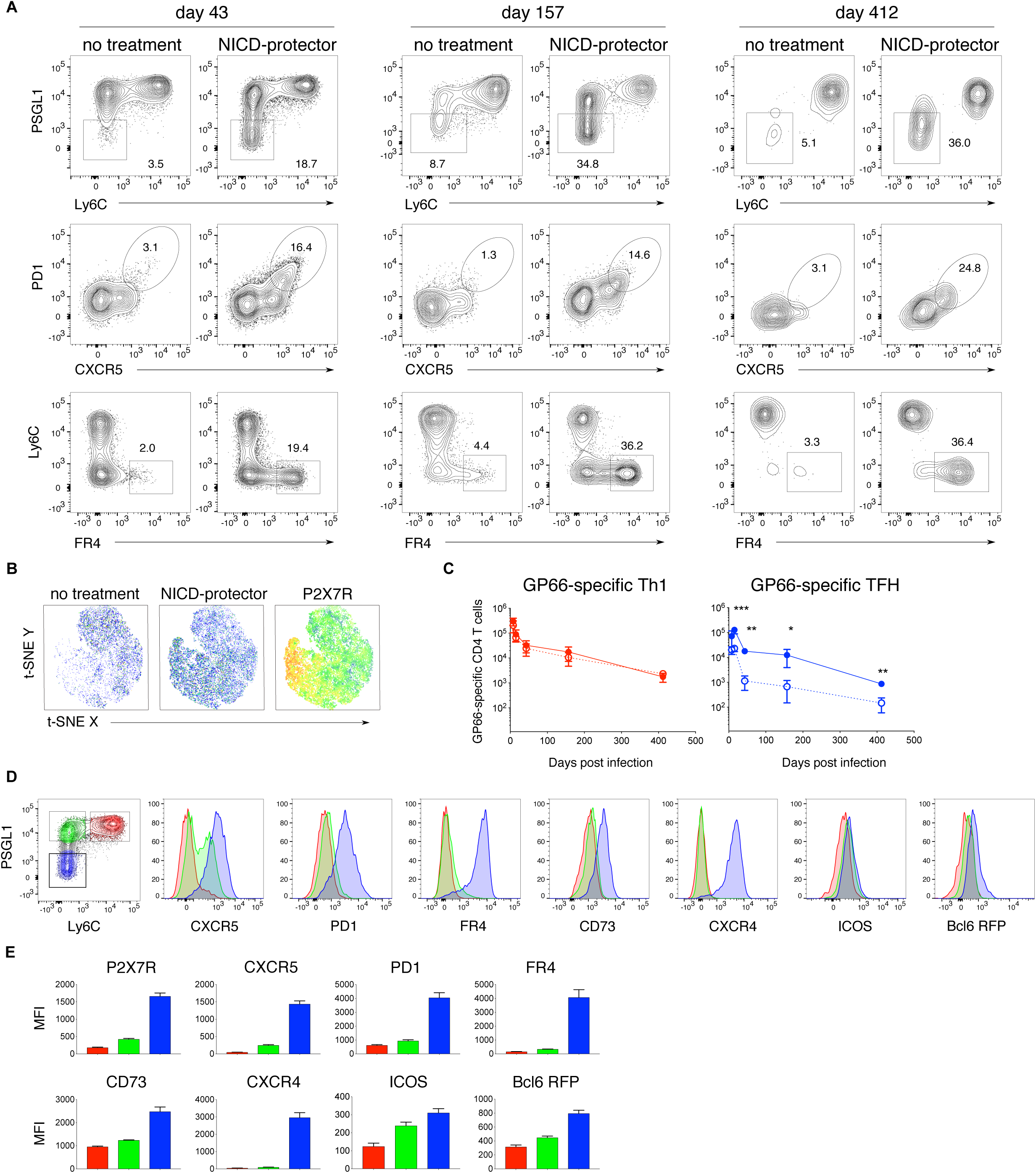
TFH memory cells are susceptible to death during isolation. (A) Flow cytometry analysis of GP66-specific CD4 T cells isolated from the spleen at the indicated time points post infection, with or without NICD-protector using different gating strategies to identify TFH memory cells (Ly6C^lo^PSGL1^lo^ or CXCR5^hi^PD1^hi^ or FR4^hi^Ly6C^lo^). (B) tSNE plots of the GP66-specific CD4 memory compartment with (middle, right) or without NICD-protector (left) and overlaid P2X7R expression with the red color indicating the highest expression level (right). (C) Quantification of GP66-specific Th1 (red, Ly6C^hi^PSGL1^hi^) or TFH memory cells (blue, Ly6C^lo^PSGL1^lo^) numbers over time with (solid line) or without (dashed line) NICD-protector gated as in Figure 1D. Thin lines represent the mean ± s.d. (D-E), Representative flow cytometry plots (D) and mean fluorescence intensity (MFI) (E) of the indicated marker in GP66-specific CD4 memory cell subsets Th1 (red), Ly6C^lo^ Th1 (green), and TFH (blue) >40 days post infection. Data represent (*N*) = 2 independent experiments for D-E with *n* = 3-4 mice per group. Unpaired two-tailed Student’s t test was performed on each individual time point (C). \**P* < 0.05, \*\**P* < 0.01, \*\*\**P* < 0.001.

### TFH memory cells are transcriptionally distinct from TCM

To gain further insight into CD4 memory T cell heterogeneity and regulation we performed single-cell RNA sequencing (scRNA-seq) on GP66-specific T cells isolated at day >35 post-infection. Principal component analysis (PCA) was used to provide a low dimensional representation of the data and as a basis for hierarchical clustering of the cells and tSNE visualization (Figures 2A and S2B). Seven distinct clusters were enriched for genes associated with: TFH cells (clusters 1-3, 37% of cells), TCM cells (clusters 4 and 5, 45% of cells) and Th1 cells (clusters 6 and 7, 18% of cells) (Figures 2A, 2B and S2B). The top defining genes in the TFH clusters included established TFH markers such as *Izumo1r* (encoding FR4), *Pdcd1, Sh2d1a*, as well as transcription factors known to be expressed by CD4 memory cells, including cells *Klf6, Jun and Junb* (Figures 2B and 2C) (Crawford et al., 2014; Iyer et al., 2013). TCM and Th1 clusters (4-7) exhibited higher expression of *Il7r, S100a4, S100a6, Selplg* and various integrins (Figures 2B and 2D), while clusters 6 and 7 were enriched for genes strongly associated with Th1 differentiation including *Cxcr6, Ccl5, Nkg7* and *Id2* (Figures 2B and 2E). Cutting the clustering tree to make only two clusters illustrates that TCM and Th memory cells are more similar to each other than either is to TFH (Figure 2B). Among Th1 clusters, cluster 7 represented the subset highest in *Selplg*, *Ly6c2* and *Il7r*, while cluster 6 expressed higher levels of *Cxcr6* and *Id2* with signatures of dysfunction and exhaustion (Figures S2D-S2F) (Ciucci et al., 2019; Martinez et al., 2015). Cluster cell-type identities were further confirmed by scoring each cell for signature genes obtained from publicly available datasets and examining the distributions of these scores across clusters (Figures 2F and S2C). TFH and Th1 signatures matched well with clusters 1-3 and 6-7 respectively, while clusters 4, 5 and 7 were enriched for the TCM precursor signature reported by Ciucci et. al. (Figure 2F) (Ciucci et al., 2019). Within TFH clusters 1-3, we observed a gradient in *Cxcr5* and *Pdcd1* expression while *Izumo1r* remained stable, serving as a much cleaner transcriptional marker of the boundary between TFH memory and TCM cells (Figures 2G and 2I). Unexpectedly, *Ccr7* expression, highest in clusters 4 and 5, was strongly negatively correlated with *Cxcr5* in cells from animals treated with NICD protector (in the bottom 4% of all genes), whereas cells from untreated animals showed a slight positive correlation (in the top 27% of all genes) (Figure 2G). This trend was mainly driven by the bona fide TFH clusters (1-3) preferentially rescued from NICD, as illustrated by the imputed expression levels of *Cxcr5* vs. *Ccr7* (Figure 2H). A similar negative correlation was found between protein expression levels for CCR7 and CXCR5, with the highest CCR7 expressing cells falling within the Ly6C^lo^ Th1 compartment (Figures 2J and S2H). Cxcr5 expression is still found in non-TFH clusters, though with a slight decrease from TCM to Th1 (Figure 2I). The use of FR4 as a marker for TFH identity thus improves upon existing strategies, which either allocate too many TCM cells into a TFH gate or depend upon markers like PD1 and CXCR5 which are known to decline over time.

**Figure 2.**
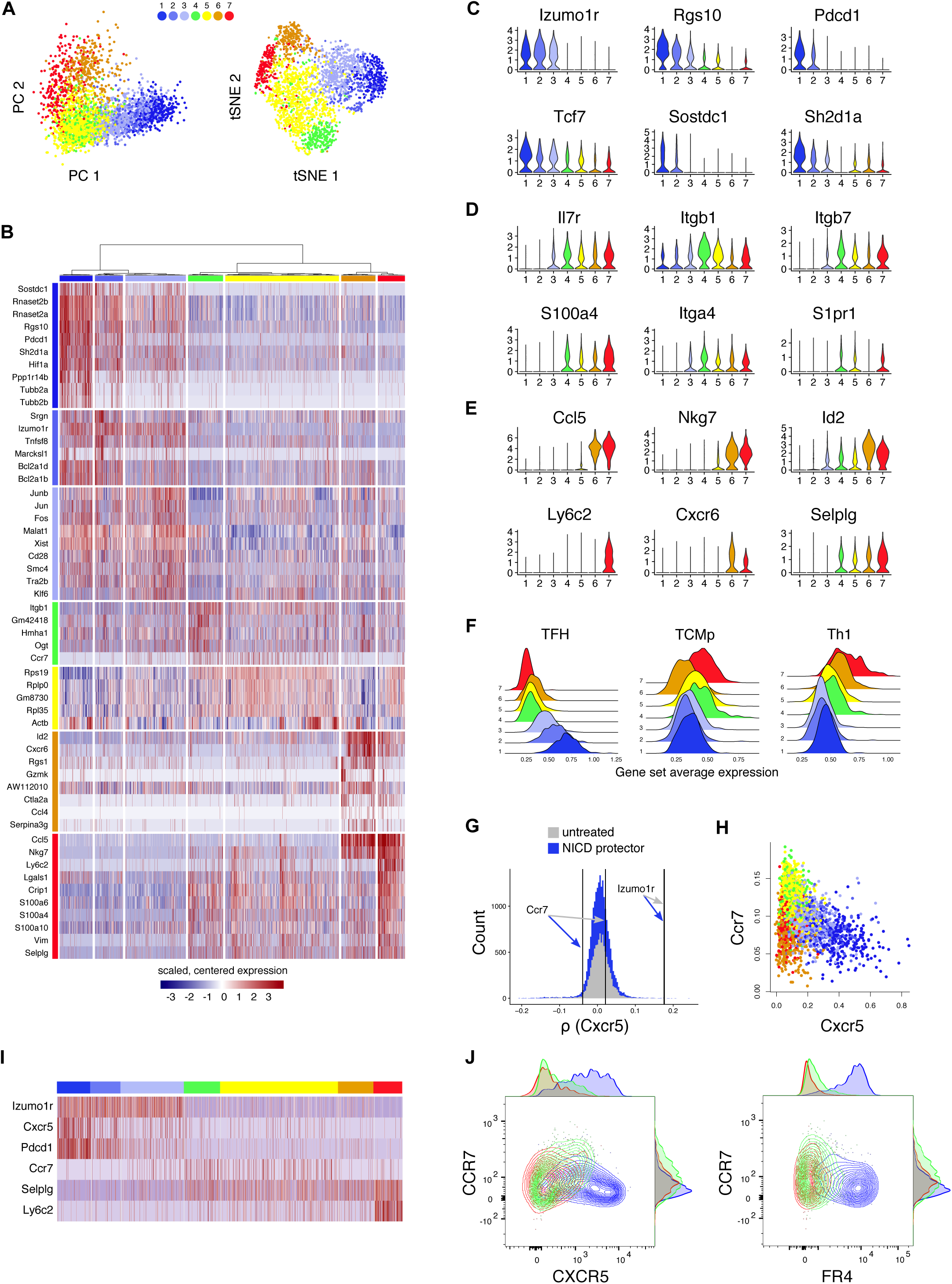
TFH memory cells are transcriptionally distinct from TCM. (A) Unsupervised hierarchical clustering using Ward’s method: PCA and tSNE dimension reductions. (B) Heatmap showing scaled, centered single cell expression data of top 10 genes according to cluster average log FC, adjusted P value < 0.05. (C-E) Log-normalized expression of genes typical for TFH (C), TCM (D) and Th1 (E) populations. (F) Log-normalized average expression of published CD4 signatures obtained by analysis of d7 effectors: TFH, TCM precursors, and Th1. (G) scRNA-seq: Rank of Ccr7 and Izumo1r (encoding FR4) among Spearman’s rank correlation coefficients between Cxcr5 and all other genes, NICD protector vs. untreated. (H) scRNA-seq: Cxcr5 vs Ccr7 on imputed data, colored by cluster. Dropout imputed using Seurat::AddImputedScore. (I) Scaled, centered single cell expression of Izumo1r in comparison to other common gating markers. Unless otherwise noted, scRNA-seq data are shown from the run with NICD-protector to ensure consistency with other experiments. (J) Flow cytometric analysis of CCR7, CXCR5 and FR4 on GP-66 specific CD4 memory subsets >50 days post infection gated as in Figure 1D. Data represent (*N)* = 2 independents with *n* = 4 mice (J).

We additionally noted that TFH memory cells expressed a partially overlapping transcriptional signature with tissue resident memory cells, emphasizing the non-circulating status of this population (Figure S2G). Notably, *Hif1a*, a transcription factor normally associated with glycolysis, was one of the top genes expressed by TFH memory cells, consistent with the recently reported high expression of this transcription factor in TFH effectors and resident memory cells (Figure 2B) (Cho et al., 2019; Hombrink et al., 2016; Zhu et al., 2019). Taken together these data highlight the clear transcriptional distinctions between ultimately more persistent NICD-rescued TFH memory and TCM cells.

### TFH memory cells are constitutively glycolytic

In addition to *Hif1a* expression, TFH memory cells were enriched for mTOR and cAMP regulated genes, suggesting a metabolic signature similar to that observed in trained immune cells (Figures 3A, S3A and S3B) (Bekkering et al., 2018; Cheng et al., 2014; Saeed et al., 2014). Enhanced activation of mTOR regulated genes was also observed following secondary analysis of microarray data on bulk sorted Smarta TFH memory cells as well as recently published single cell data from GP66-specific TFH memory cells (Figures S3C and S3D), and was further confirmed by qPCR of selected Raptor regulated genes in sorted CD4 GP66-specific memory T cell populations (Figure 3B) (Zeng et al., 2016). These data indicate that TFH memory cells may constitutively engage a glycolytic metabolism. To assess mTORC1 activity, we measured phosphorylation of TORC1-target ribosomal protein S6 (p-S6) directly after T cell isolation (Figures 3C and S3E). Both GP66-specific TFH cells as well as CD44+ TFH cells, which have phenotypic characteristics similar to antigen-specific TFH cells (Figure S3F), exhibited increased p-S6 compared to Th1 memory cells (Figures 3C and S3E). Consistent with mTOR activation, TFH memory cells had increased uptake of the fluorescent glucose analog 2-NBDG, and an increased baseline extracellular acidification rate (ECAR) (Figures 3D and 3E). However, alongside this apparent increase in glycolytic metabolism, TFH memory cells had slightly decreased expression of the amino acid transporter CD98 and the transcription factor c-myc compared to Th1 memory cells, indicating that not all measures of anabolic activity are increased in these cells (Figures S3G and S3H).

**Figure 3.**
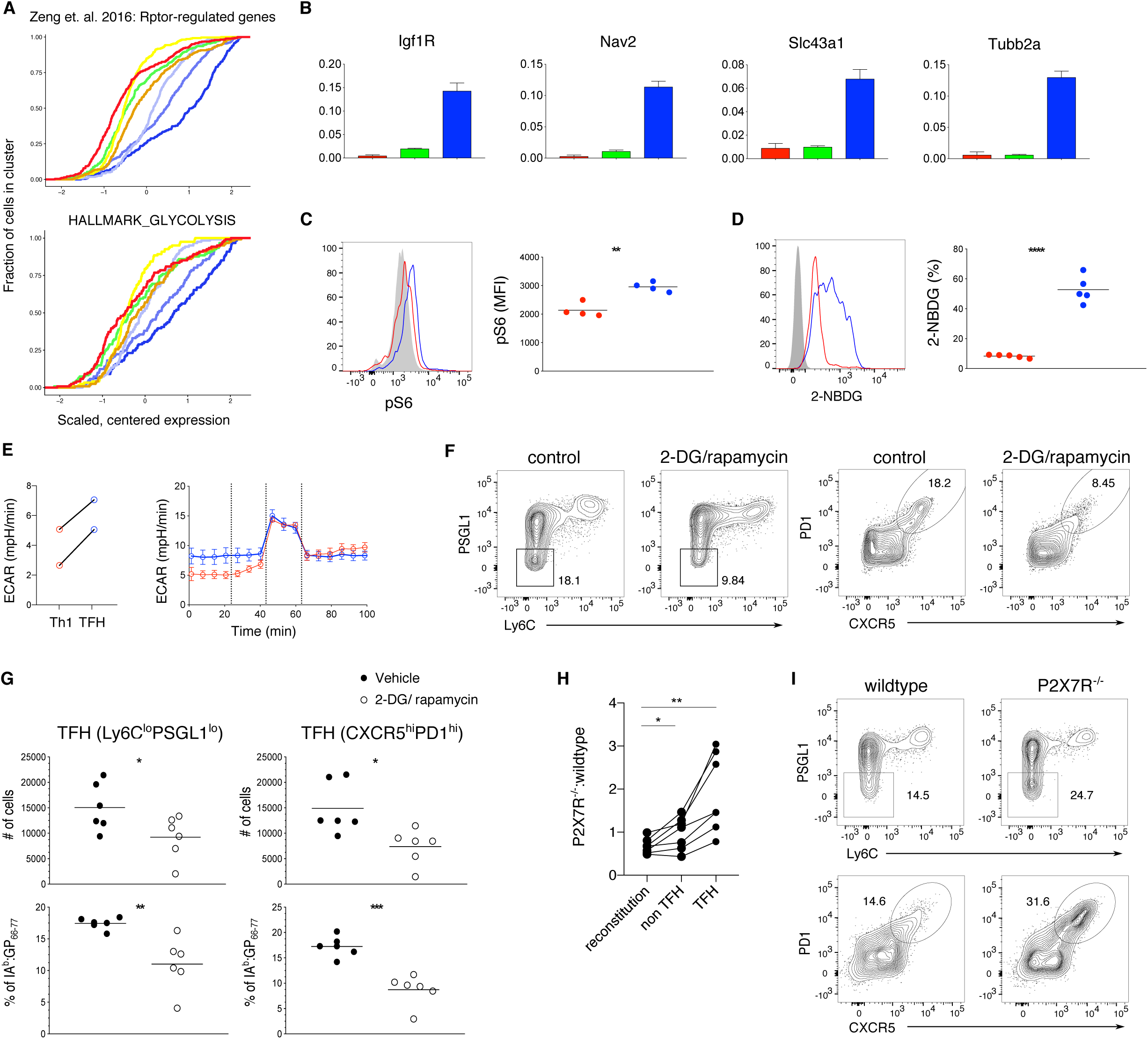
TFH memory cells are constitutively glycolytic. (A) Cumulative distribution of scaled, centered mRNA expression by cluster of Rptor-regulated genes (Zeng et al., 2016) and the gene set HALLMARK_GLYCOLYSIS. (B) quantitative PCR of genes (arbitrary units relative to Rpl13a) in sort-purified GP66-specific Th1 (red), Ly6C^lo^ Th1 (green) and TFH (blue) memory cells gated as in Figure 1D. (C-D) Representative histograms and MFI of intracellular phospho-S6 Ser240/244 on CD44+ Th1 or TFH memory cells compared to naive CD4 T cells (grey) (C) and 2-NBDG uptake (D) on GP66-specific memory cells >30 days post infection compared to FMO control (grey). (E) Quantification of basal ECAR and ECAR profile in response to Mito-Stress test in Th1 and TFH memory cells pooled from 20-30 mice. (F-G) Representative flow cytometry plots (F) and quantification (G) of GP66-specific memory cells treated with vehicle or 2-DG/rapamycin. (H-I) P2X7R^−/−^ to wildtype ratio of GP66-specific memory TFH cells >60 days post infection (H) and analysis (I) using different gating strategies. Data is representative of *(N)*= 2 independent experiments depicting the mean ±s.d. of *n* = 3 technical replicates (B) or 4 (C), 5 (D), 6 (F-G) or 7 (H-I) mice per group. Dots represent cells from individual mice, and the line represents the mean (C, D, G). Unpaired (C, D, G) or paired (H) two-tailed Student’s t test was performed with \**P* < 0.05, \*\**P* < 0.01, \*\*\**P* < 0.001, \*\*\*\**P* < 0.0001.

To determine if the shift toward glycolysis by TFH memory cells was accompanied by a reduction in oxidative metabolism we also assessed their baseline oxygen consumption rate (OCR). Although we observed a slight reduction in OCR by TFH memory cells compared to Th1 memory cells, the maximum respiration and spare respiratory capacity were similar between these two populations (Figure S3I). At the transcriptome level, TFH memory cells were also enriched for genes associated with oxidative phosphorylation and mitochondrial respiration, a signature commonly associated with CD8 memory T cell survival (Figure S3J). As TFH effectors were previously reported to have decreased mTOR activation and reduced glycolysis (Yang et al., 2016; Zeng et al., 2016), we also assessed the metabolic capacity of these cells isolated at day 10 after infection, reasoning that NICD-protector might improve their survival and fitness. At this time point, TFH effectors had slightly reduced basal ECAR compared to Th1 effectors but demonstrated normal glycolytic flux following the addition of FCCP (Figure S3K). In addition, and similar to TFH memory cells, TFH effectors had equivalent baseline and maximal respiratory capacity as Th1 effectors indicating that in contrast to an earlier report, NICD-protector also preserves the metabolic fitness of TFH cells isolated at earlier time points after infection (Figure S3L) (Marshall et al., 2011). To assess whether constitutive glycolysis/mTOR signaling are required to maintain long lived TFH cells, we next examined TFH cell survival in mice treated with the glycolysis inhibitor 2-deoxyglucose (2-DG) and rapamycin starting at day >40 after LCMV infection. After two weeks of 2-DG/rapamycin treatment, both the proportion and number of TFH memory cells were significantly decreased, with the remaining TFH memory cells showing a reduction in size consistent with reduced mTOR activation (Figures 3F, 3G and S3M) (Ray et al., 2015). Signaling through P2X7R was recently suggested to restrain mTOR activation, leading to improved survival of CD8 memory T cells (Borges da Silva et al., 2018). In contrast, P2X7R signals have been shown to restrain accumulation of TFH effector cells in Peyer’s Patches (Proietti et al., 2014). To assess whether P2X7R signals promote or restrain TFH memory cell survival, we generated radiation bone marrow chimeras with a mixture of wild-type and P2X7R-deficient bone marrow. Sixty days after LCMV infection, P2X7R-deficient T cells generated an increased proportion of all memory cells, with the most significant increase in the TFH memory compartment (Figures 3H and 3I). These data demonstrate that P2X7R is a key negative regulator of TFH memory cell accumulation and are consistent with a role for P2X7R in restraining mTOR activation.

### TFH memory cells can survive in the absence of antigen

Glycolytic metabolism in T cells is often associated with activation and effector cell proliferation. To understand if TFH memory cells are responding to ongoing antigen stimulation we examined Nur77 expression within the TFH memory compartment of LCMV infected Nur77 reporter mice. Approximately 30% of PSGL1^lo^Ly6C^lo^ TFH cells were Nur77 positive, with a slightly higher percentage in the CXCR5^hi^PD1^hi^ TFH compartment (Figure 4A). While PD1 expression was moderately increased on Nur77+ compared to Nur77-TFH memory cells, it was much higher on both of these populations compared to non-TFH memory cells (Figure 4B). Although these data indicate that a fraction of the TFH memory compartment continues to respond to antigen, we did not observe a positive correlation between PD1 expression and 2-NBDG uptake, indicating that glucose uptake within the TFH compartment is unlikely to be related to antigen stimulation (Figure S4A). Consistent with this idea, increased 2-NBDG uptake was maintained by TFH memory cells at >400 days of infection, a time point where PD1 expression by TFH memory cells is decreased compared to earlier time points (Figure S4B). To understand whether the glycolytic metabolism observed in TFH memory cells is related to proliferation, we administered bromo-2-deoxyuridine (BrdU) in the drinking water of LCMV infected mice, beginning 40 days after infection. After 12-14 days, approximately 4% of PSGL1^lo^Ly6C^lo^ TFH memory cells incorporated BrdU with a slightly higher proportion CXCR5^hi^PD1^hi^ TFH cells staining positive for BrdU (Figure 4C). Ly6C^lo^ Th1 memory cells showed a similar level of in vivo cycling with approximately 5% of cells staining for BrdU, while 15% of Ly6C^hi^ Th1 memory cells were BrdU positive, demonstrating more extensive proliferation by this subset (Figure 4C). These data show that the glycolytic phenotype observed in TFH memory cells is not strictly correlated with extensive cell division, although the slightly increased BrdU incorporation by CXCR5^hi^PD1^hi^ TFH memory cells may be related to ongoing antigen stimulation. To better understand the contribution of antigen to maintaining the TFH memory compartment, we transferred LCMV-specific effector cells isolated at day 10 after infection into either LCMV infection matched or naïve recipients followed by analysis of donor T cell phenotype. Thirty days after primary infection, Ly6C^hi^ Th1 memory cells were nearly absent in both transfer scenarios, indicating that Th1 effector cells are susceptible to death at the time point of isolation and transfer (Figure 4D). In contrast, GP66-specfic TFH memory cells were detected in both antigen free (naïve) and infection matched mice. Although lower numbers of TFH memory cells with reduced expression of CXCR5 and PD1 were recovered from naïve mice, this may be partly due to ongoing expansion of TFH effector cells in LCMV infected recipients between days 10-15 after transfer (Figures 4D, 4E and 1C). Nevertheless, these TFH memory cells maintained high expression of FR4, consistent with scRNA-seq data pinpointing FR4 (encoded by *Izumo1r*) as a more reliable marker for bona fide TFH memory cells (Figures 4D, 4E, 2C and 2I). Taken together, these data underscore the ability of TFH memory cells to survive in the absence of antigen.

**Figure 4.**
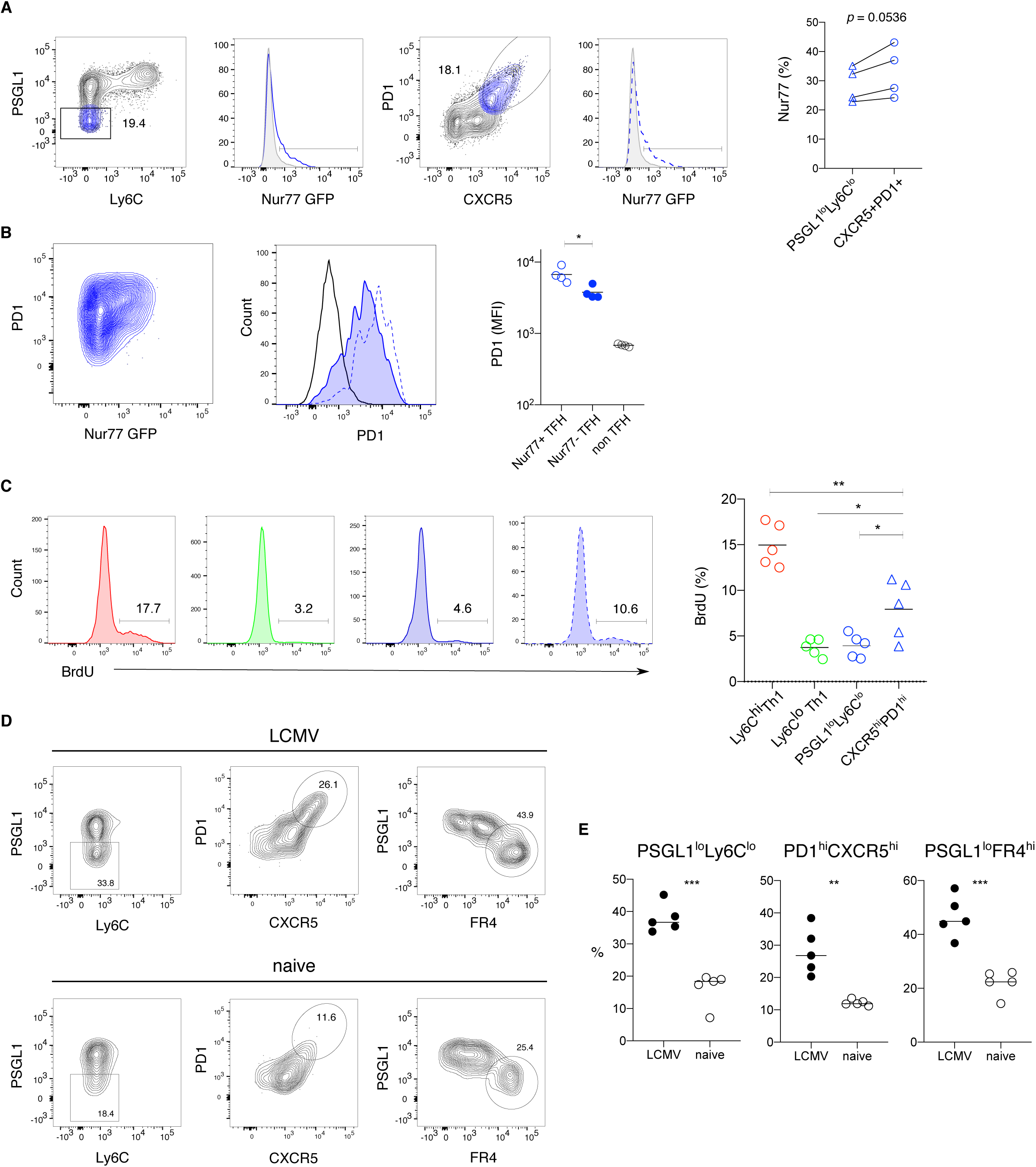
TFH memory cells can survive in the absence of antigen. (A) Flow cytometry analysis of Nur77 expression by GP66-specific TFH memory cells for the indicated gating strategies. (B) Representative flow cytometry plots (left panels) and MFI (right panel) depicting PD1 expression on Nur77+ (dotted blue line) and Nur77-(tinted blue line) TFH cells, gated as depicted in panel (A). (C) Flow cytometry plots (left) and quantification (right) of BrdU incorporation by GP66-specific CD4 memory subsets after providing BrdU for 12 days in drinking water. (D-E) Total GP66-specific CD4 T cells were isolated at day 10 after LCMV infection and transferred into congenic infection matched (LCMV) or naive mice. Donor cell phenotype was analyzed at day 30 after infection. Representative flow cytometry plots (D) and quantification (E) with gating for PSGL1^lo^Ly6C^lo^ (left), PD1^hi^CXCR5^hi^ (middle) and PSGL1^lo^FR4^hi^ (right) with dots representing cells from individual mice, and the line depicts the mean. Data represent *(N)* = 2 independent experiments with n = 4-5 mice. Paired (A) or unpaired two-tailed Student’s t test (B,C,E) was performed with \**P* < 0.05, \*\**P* < 0.01, \*\*\**P* < 0.001.

### TFH memory cells generate multiple cell fates upon recall

Despite maintaining a glycolytic phenotype, TFH memory cells are also enriched for genes regulated by *Tcf7* (encoding TCF1) and highly express *Cd27*, a marker associated with memory T cell survival, cytokine production potential and stemness (Figures 5A and S5A) (Hendriks et al., 2000; Karmaus et al., 2019; Pepper et al., 2010). Although higher expression of Tcf7 in TFH memory cells translated into higher expression of TCF1 protein, CD27 protein expression was significantly lower in TFH compared to Th1 memory cells (Figures 5B and S5B). To determine if the discrepancy between the CD27 gene expression and protein expression was a result of P2X7R mediated shedding, we examined CD27 on memory cells isolated from LCMV infected mixed bone marrow chimeras reconstituted with an equal mixture of P2X7R-deficient and wild type bone marrow cells (Moon et al., 2006). Here we observed higher expression of CD27 on P2X7R-deficient TFH memory cells compared to both wild type TFH and Th1 memory cells (Figure S5B). Consistent with high expression of both *Tcf7* and *Cd27*, TFH memory cells were also enriched for “early memory” associated genes (Figure S5C). In addition, reconstruction of a cell developmental trajectory indicated that TFH and Th1 memory cells occupied opposite ends of a pseudotime trajectory, predicting enhanced differentiation plasticity in one of these subsets (Figures S5D and S5E) (Trapnell et al., 2014). To test this in vivo, TFH (FR4^hi^PSGL1^lo^Ly6^lo^) and Th1 (FR4^lo^PSGL1^lo^Ly6^hi^ and FR4^lo^PSGL1^hi^Ly6^hi^) memory cells were sorted from LCMV infected mice and transferred into naïve congenic recipients, followed by secondary challenge with LCMV (Figure 5D). Twelve days after recall infection, transferred TFH memory cells differentiated into both TFH and Th1 effectors, demonstrating lineage flexibility within this subset (Figures 5E and 5F). In contrast, Ly6C^lo^ Th1 memory cells maintained the capacity to differentiate into both Ly6C^hi^ and Ly6C^lo^ effectors, but gave rise to very few TFH effectors, suggesting more limited plasticity of this subset (Figures 5E and 5F). Consistent with several previous reports, Ly6C^hi^ Th1 memory cells almost exclusively gave rise to Ly6C^hi^ secondary effectors, indicating more terminal differentiation of these cells (Figures 5E and 5F) (Hale et al., 2013; Pepper et al., 2011). Taken together, these data demonstrate that despite engaging an anabolic metabolism often associated with effector cell proliferation and differentiation, TFH memory cells maintain the capacity to differentiate into multiple types of effectors following secondary challenge.

**Figure 5.**
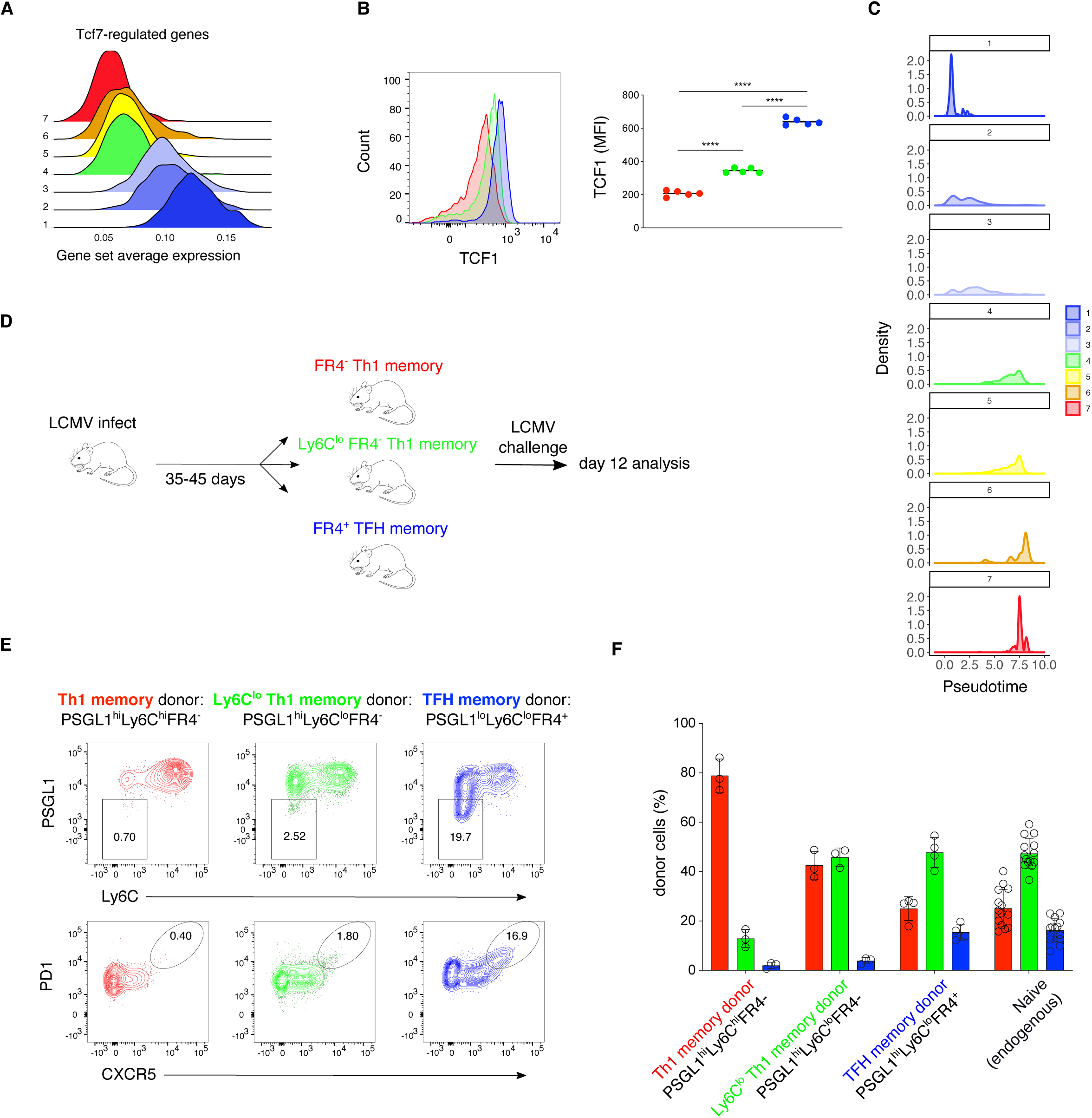
TFH memory cells generate multiple cell fates upon recall. (A) Log-normalized average expression of genes tracking with Tcf7 (Utzschneider et al., 2016). (B) Flow cytometry analysis of TCF1(left) and MFI (right) in Th1 (red), Ly6C^lo^ Th1 (green) and TFH (blue) GP66-specific memory cells. (C) Density of cells in pseudotime, Monocle2 analysis. (D) GP66-specific Ly6C^hi^ Th1, Ly6C^lo^ Th1 or TFH CD4 memory T cells were sorted at day 35-45 after LCMV infection and transferred into congenic naive mice. The recipient mice were challenged with LCMV the following day. Donor cell phenotype was analyzed at day 12 after infection. (E) Representative plots of flow cytometric analysis of Ly6C^hi^ Th1 (red), Ly6C^lo^ Th1 (green) or TFH (blue) donor cell phenotype. (F) Quantification of Ly6C^hi^ Th1 effectors (red), Ly6C^lo^ Th1(green) and TFH (blue) effectors from Ly6C^hi^ Th1, Ly6C^lo^ Th1 or TFH donors compared to endogenous response. Data represents *(N)* = 2 independent experiments with *n* = 5 (B) or 3-14 (E-F) mice per group. One-way ANOVA followed by Turkey’s post-test for multiple comparisons with \*\*\*\**P* < 0.0001.

### Epigenetic regulation of TFH memory T cells

To assess whether the distinct transcriptional signatures observed in TFH and Th1 memory cells were also apparent at the epigenetic level we compared chromatin accessibility of naïve, Th1 and TFH memory T cells by ATAC-seq (Buenrostro et al., 2013). The majority of called peaks were shared among these 3 populations, with the highest number of unique peaks present in naïve CD4 T cells (Figure S6A). Principal component analysis (PCA) revealed that TFH and Th1 memory cells are set apart from naïve T cells along PC1, yet separated from each other along PC2, suggesting they possess distinct accessibility profiles (Figure 6A). While Th1 memory cells had more differentially open peaks annotated to exons, introns and distal intergenic regions, TFH memory cells had about 50% more differentially accessible peaks located in promoter regions (Figure S6B). We next focused on the significantly differentially accessible promoter regions with a high fold change (log_2_ fold change > 1 and false discovery rate < 0.05) between at least two of the T cell subsets and performed hierarchical clustering of genes and samples. This resulted in clusters of genes with common accessibility patterns: e.g. up in naïve, up in memory, up in naïve and TFH memory (Figures 6B and S6C). For example, the *Foxp1* promoter was exclusively accessible in naïve CD4 T cells, consistent with its role as a gatekeeper of the naïve to memory T cell transition (Durek et al., 2016). Integrative analysis of ATAC-seq and RNA-seq data (*in silico* pseudo-bulk samples derived from scRNA-seq data) revealed a correlation between the promoter accessibility and expression of genes defining TFH and Th1 memory cell subsets (Figure S6D) (Hale et al., 2013). Consistent with a study that assessed the DNA methylation status of various cytokine loci in TFH and Th1 memory cells (Hale et al., 2013), we observed that the *Gzmb* promoter was exclusively accessible in Th1 memory cells, while the *Ifng* promoter was accessible in both populations, albeit at slightly lower levels in TFH memory cells (Figure S6E). In contrast to this earlier study, however, the promoter accessibility of the hallmark TFH cytokine, *Il21*, was restricted to TFH memory cells (Figure S6E). In addition, one of the most accessible cytokine promoter regions in TFH memory cells was the epidermal growth factor-like ligand amphiregulin (*Areg*), notable for its role in restoring tissue integrity after infection (Zaiss et al., 2015). To further assess differences in the transcriptional regulation of TFH and Th1 memory cells, we analyzed enriched binding motifs in called peaks using Hypergeometric Optimization of Motif Enrichment (HOMER) (Heinz et al., 2010). In agreement with the RNA-seq data, TFH memory cells were enriched for motifs belonging to the Tcf7 transcription factor family as well as AP-1 family members which are known to be similarly regulated in CD8 memory cells (Figure 6C) (Lau et al., 2018; Moskowitz et al., 2017). In contrast, Th1 memory peaks exhibited increased Runx and T-box transcription family member binding sites, consistent with the cooperation of these transcription factors in regulating IFNγ and stabilizing Th1 fate in CD4 T cells (Figure S6F) (Djuretic et al., 2007; Kohu et al., 2009). These observations were further validated by applying the single-cell regulatory network interference and clustering (SCENIC) algorithm to scRNA-seq data, which highlighted Tcf7/AP-1 and Runx family members as regulators in TFH memory and Th1 memory, respectively (Figure S6G). We next assessed specific functional pathways in the ATAC-seq data by running gene set enrichment analysis (GSEA) on differentially accessible promoter regions in TFH compared to Th1 memory cells using both standard gene set categories as well as sets curated from relevant publications (Figures 6D and S6H). Here we observed increased promoter accessibility in TFH memory cells of *Rptor* and *Tcf7* regulated genes as well as genes activated in response to cAMP in TFH memory cells, suggesting that the anabolic and stem signatures of this population are strongly regulated at the epigenetic level (Figures 6D and S6H). Finally, using Ingenuity Pathway Analysis (IPA) (Qiagen, version 01-14) to probe the functional regulation of TFH memory cells, we identified ICOS-ICOSL signaling as one of the top pathways enriched in both ATAC-seq and scRNA-seq data sets (Figure 6E) (Kramer et al., 2014). These data raise the possibility that ICOS mediated signals contribute to the maintenance and identity of TFH memory via specific epigenetic modifications.

**Figure 6.**
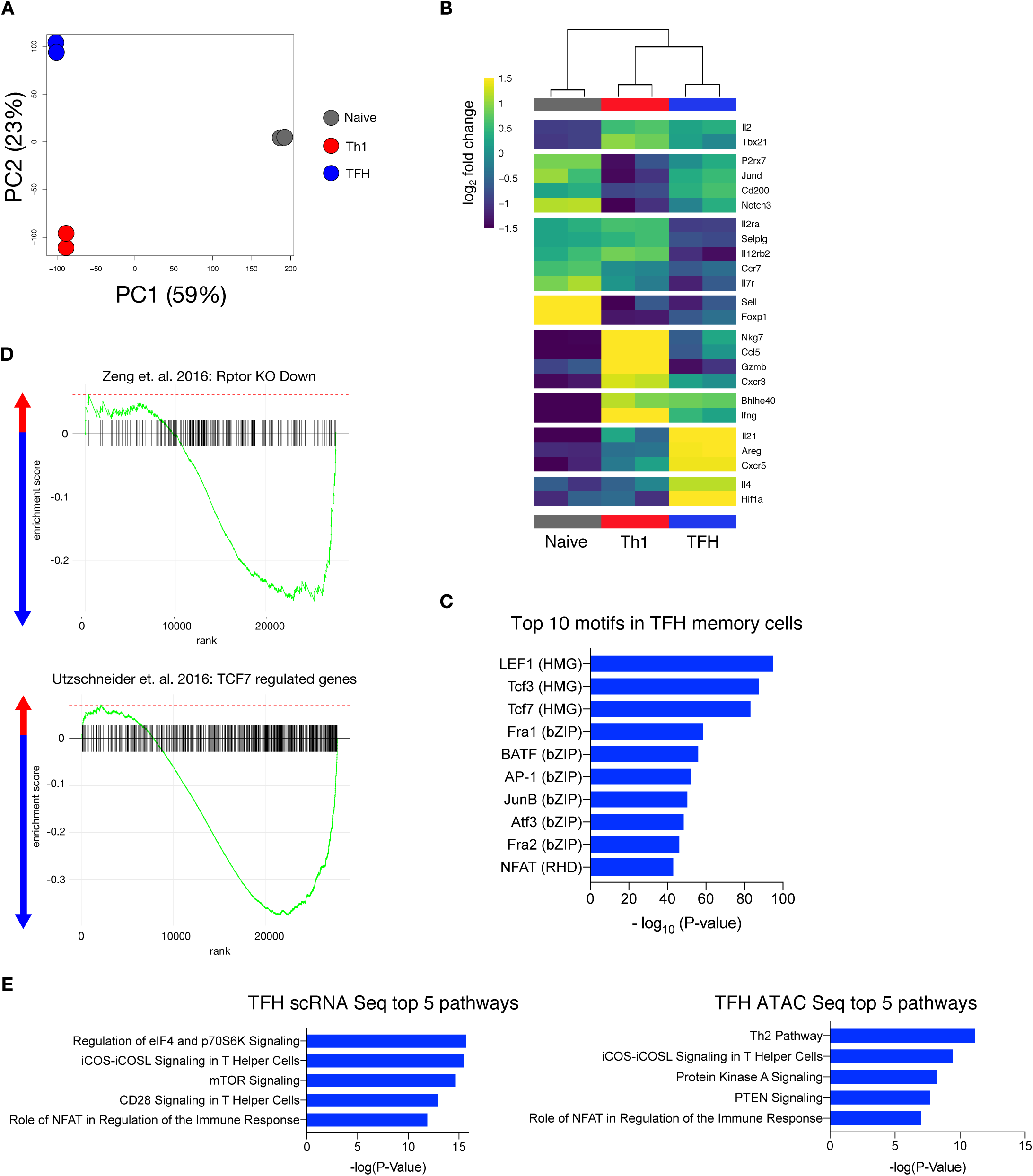
Epigenetic regulation of TFH memory cells. (A) PCA of Assay for Transposase-Accessible Chromatin using sequencing (ATAC-seq) data on all peaks. (B) Heatmap of hierarchically clustered promoter regions of highlighted genes. Genes with absolute value (log_2_ fold change) > 1 are assigned the most extreme colors. (C) Enriched motifs in TFH memory cells found by HOMER. (D) GSEA analysis of Rptor (above) and Tcf7 (below) regulated genes with negative enrichment score indicating enrichment in TFH memory and positive score indicating enrichment in Th1 memory. (E) Top 5 pathways enriched in TFH memory cells determined by Ingenuity Pathway Analysis in: combined scRNA-seq data (with and without NICD-protector) comparing TFH cluster to Th1 cluster from 3-cluster version (left) and ATAC-seq peaks mapped to nearest TSS within TAD (right).

### ICOS signaling maintains TFH memory cell identity

Although ICOS has an established role in primary TFH responses it’s role in maintaining CD4 memory is less clear. ICOS signaling has been shown to induce the expression of Tcf7 in Th17 memory cells that maintain the plasticity to differentiate following stimulation (Majchrzak et al., 2017). ICOS can also induce mTORC1 activation in both TFH effector and T follicular regulatory (TFR) effector cells, leading to further stabilization of the follicular T cell program (Xu et al., 2017; Yang et al., 2016; Zeng et al., 2016). To test the hypothesis that ICOS signals contribute to the integration of stemness and anabolic metabolism in TFH memory cells we treated LCMV infected mice with anti-ICOSL starting at day >50 after infection (Figure 7A). After 12 days of treatment, the number of GP66-specific TFH cells was decreased in anti-ICOSL treated mice (Figure 7B). The TFH memory cells remaining after ICOS blockade had decreased expression of TCF1 as well as reduced activation of mTOR as measured by cell size (Figures S7A and S7B). To further characterize the effects of anti-ICOSL blocking on TFH memory cells, we performed RNA-seq at day 60 post infection on control and treated mice. Cells from mice treated with anti-ICOSL for 12 days prior to harvest exhibited lower transcription of key TFH genes such as *Pdcd1*, *Cxcr5* and *Bcl6*, while known Th1 survival factors such as *Bcl2* and *Il7r* were increased (Figure 7C). Gene set enrichment analysis showed decline of TFH-defining, resident memory, *Rptor*-regulated and stem cell-associated sets in treated samples compared to controls (Figure 7D). These data emphasize the importance of ICOS signals in maintaining TFH memory cell identity.

**Figure 7.**
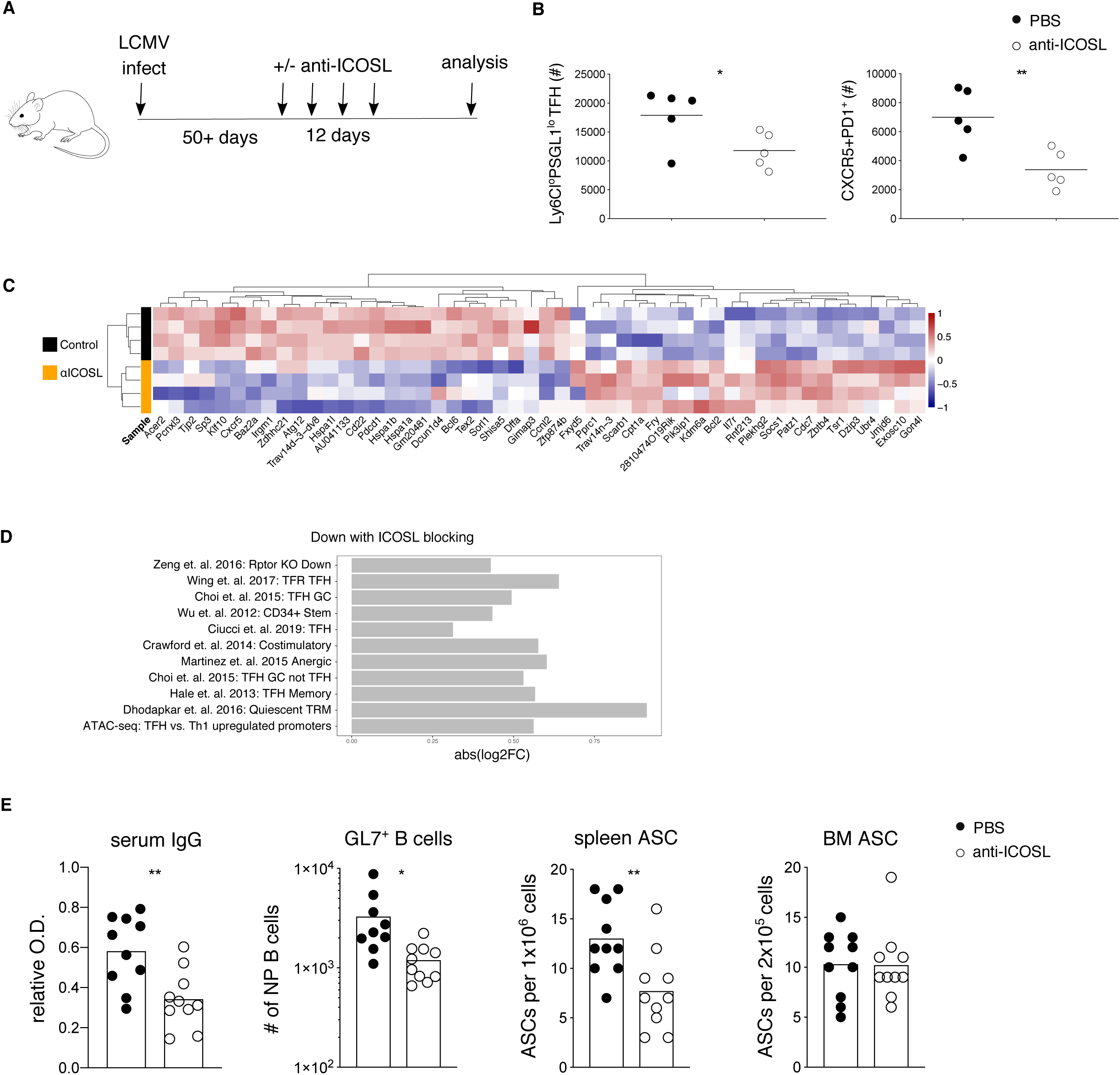
ICOS signaling maintains TFH memory cell identity. (A) Mice were infected with LCMV followed by anti-ICOSL treatment at late time points after infection. LCMV specific CD4 T cells and B cells were analyzed after 12 days of treatment. (B) Cell numbers of GP66-specific TFH memory cells in treated mice using either Ly6C^lo^PSGL1^lo^ or CXCR5^hi^PD1^hi^ as gating strategy. (C) Scaled, centered expression of 20 most significant differentially expressed transcripts in TFH memory cells between control and anti-ICOSL treated mice. (D) Most significantly downregulated gene signatures with anti-ICOSL blocking (FDR < 0.2) (Boddupalli et al., 2016; Choi et al., 2015; Ciucci et al., 2019; Crawford et al., 2014; Hale et al., 2013; Martinez et al., 2015; Wing et al., 2017; Wu et al., 2012; Zeng et al., 2016). (E) NP-specific IgG serum titer (far left), numbers of NP-specific GL7+ B cells (left) and NP-specific antibody secreting cells from the spleen (right) and bone marrow (far right). Data represents one of *(N)* = 2 independent experiments with *n* = 5 mice per group (B). Data summarize (*N)* = 2 independent experiments with *n* = 9-10 mice per group and bars indicating the mean (E). Unpaired two-tailed Student’s t test was performed for (B, E) with \**P* < 0.05, \*\**P* < 0.01.

During primary immune responses, ICOS signals promote TFH effector cell localization in B cell follicles and primary germinal center responses (Shi et al., 2018; Weber et al., 2015). To assess whether late ICOS blockade and subsequent decline of the TFH memory cell compartment impacts late phase humoral responses, we measured LCMV-specific antibody titers in the serum of treated and control mice at day >60. Here we observed an almost 2-fold decrease in NP-specific antibody titers (Figure 7E). At this late time point after infection circulating virus-specific antibodies are maintained by plasma cells, approximately 80% of which are localized in the bone marrow compartment (Slifka and Ahmed, 1996; Slifka et al., 1995). However, plasma cells are also observed in the spleen, with an estimated half-life of 172 days (Slifka et al., 1998) (Ndungu et al., 2009). To determine if the decline in antibody titers correlated with fewer plasma cells in the bone marrow or splenic compartments, we measured the number of NP-specific plasma cells by ELISPOT. Here we observed that anti-ICOSL treatment correlated with a two-fold decrease in the number of splenic plasma cells, with no apparent impact on the number of bone marrow plasma cells (Figure 7E). Assuming that ICOS blocking is 100% efficient and that circulating IgG has a half-life of approximately 8 days, these data reveal a significant and ongoing contribution of splenic plasma cells to antibody maintenance at this time point (Figure S7C). To further assess humoral immune responses, we examined virus specific B cell responses using an NP-tetramer in combination with a decoy reagent to exclude non-specific B cells (Taylor et al., 2012). Neither the total number of NP-specific memory B cells nor the number of IgD, IgM or swIg memory B cells were affected by ICOS blockade (Figure S7D). There was a significant decrease, however, in the number and proportion of GL7+ B cells (Figure S7E). Of note, these cells did not express Fas, a marker normally associated with GC B cells and that when absent, has been reported to preferentially support plasma cell differentiation (Figure S7E) (Butt et al., 2015). Taken together, these data demonstrate an essential role for ICOS signaling in maintaining the TFH memory compartment and plasma cell survival in the spleen.

## DISCUSSION

In this study we determined that the CD4 memory compartment contains an abundant and highly persistent population of TFH memory cells with broad recall capacity following secondary challenge. The presence of TFH memory has been controversial. While cells with TFH memory cell markers have been identified in the memory phase after infection, it is unclear if they represent an ongoing immune response, if they are ‘rested down’ TFH memory cells, or if they are a distinct population of self-renewing TCM cells with TFH-like characteristics (Nguyen et al., 2019). Our data demonstrating the unique susceptibility of TFH memory cells to death during isolation from the tissue shed light on the previous difficulty in discriminating these possibilities. To begin with, earlier studies reporting that TFH memory cells have a muted phenotype, with decreased or absent expression of CXCR5 and PD1, respectively, were conducted with monoclonal populations of transferred TCR transgenic cells that may not accurately reflect the breadth of a polyclonal repertoire (Hale et al., 2013; Marshall et al., 2011; Tubo et al., 2013). In addition to demonstrating more limited TFH memory differentiation by two distinct TCR transgenic strains, we show that treatment with NICD-protector rescues polyclonal TFH memory cells, preserving their phenotype and highlighting their unexpected longevity. In contrast, TCM cells, generally considered to be self-renewing and multipotent, are largely absent at the latest time points after infection. Previous studies have suggested the existence of memory cells with a hybrid TFH/TCM phenotype, namely dual expression of CXCR5 and CCR7 (Ciucci et al., 2019; Marshall et al., 2011; Pepper et al., 2011). Our data on unprotected samples confirm a population of CXCR5+CCR7+ memory cells that likely represents cells on the border of transition from TFH to TCM/Th1. However, the addition of NICD-protector reveals that these transitional cells constitute a minority of the total memory population. In addition, we show that CXCR5 as a marker to define the TFH memory compartment is complicated by broader expression of this protein in non-TFH subsets. Using scRNA-seq and flow cytometry we demonstrate that these populations are more clearly discriminated by *Izumo1r/*FR4 expression.

We further show that TFH memory cells require ongoing ICOS signals and mTOR/glycolysis for their survival. These observations suggest that maintenance of TFH memory cells is an active process. In the absence of ICOS signaling, TFH memory cells may re-localize and acquire Th1 memory cell characteristics as previously reported for TFH effector cells (Weber et al., 2015). This idea is consistent with our experiments demonstrating the inherent plasticity of TFH memory cells following transfer, although under unperturbed conditions, ICOS mediated retention of TFH memory cells in proximity with B cells may limit their secondary effector potential. These data are also in line with a recent study showing the importance of ICOS signals for the maintenance of lung resident CD4 T cells induced by tuberculosis infection (Moguche et al., 2015). Our scRNA-seq data reveal transcriptional similarity between TFH memory and resident memory T cells, both recognized as non-circulating subsets. As both residency and mTOR gene expression signatures are reduced in TFH memory cells following ICOS blockade, it will be interesting to determine if ICOS and mTOR signals are required to sustain resident memory T cells following acute infection.

The constitutive glycolytic phenotype we observed in TFH memory cells occurs together with high expression of TCF1 and CD27, both associated with self renewal and plasticity. These data are noteworthy in light of the many studies showing a correlation between catabolic metabolism and stem-like features. In general, effector T cell differentiation is associated with a more glycolytic metabolism while memory T cells exhibit a shift toward oxidative metabolism (Bantug et al., 2018). Previous studies have shown that inhibition of mTOR signals promotes the generation of both CD4 and CD8 memory T cells (Araki et al., 2009; Pearce et al., 2009; Ye et al., 2017). mTOR activation was also recently shown to mediate the differentiation of metabolically quiescent, TCF1-positive Th17 cells into inflammatory Th1 effectors (Karmaus et al., 2019). On the other hand, CD4 memory T cells require Notch dependent glucose uptake for survival, and CD8 T cells with enforced glycolytic metabolism are still able to generate memory T cells with robust recall capacity, albeit with a bias toward the CD62L^lo^ T effector memory (TEM) phenotype (Maekawa et al., 2015; Phan et al., 2016). Similarly, human CD4 TEM cells have an increased glycolytic reserve and depend on glycolysis to maintain mitochondrial membrane potential, regulating both survival and recall potential (Dimeloe et al., 2016). Taken together, our findings suggest that engagement of a specific metabolic pathway does not universally drive the differentiation of memory T cells, and that different types of memory T cells may preferentially employ metabolic pathways that are tuned to a particular immune context (e.g. availability of cytokines, nutrients, antigen, etc.).

Finally, our data reveal an unexpected role for TFH cells in supporting plasma cells and antibody titers during the memory phase, when the immune response has supposedly died down. Due to the apparent absence of TFH memory cells without NICD protector, a role for TFH cells was not previously considered. One possibility is that persistent antigen fuels the continuous differentiation of memory B cells into plasma cells (Ochsenbein et al., 2000). The observation of decreased NP-specific GL7+ B cells following ICOS blockade is compatible with this idea. However, we did not see an impact on NP-specific plasmablasts (data not shown), though we cannot exclude that this might be due to technical limitations. Another possibility is that ICOS blocking interferes with the structural organization of the spleen and/or impairment of the plasma cell niche. Plasma cell survival depends on the availability of soluble factors, such as IL-6, IL-21 and BAFF, all of which can be produced by TFH cells (Tangye, 2011). Both CD4 T cells and BAFF were recently identified as key factors specifically supporting the survival of splenic plasma cells, but the nature of CD4 derived signals was not reported (Thai et al., 2018). One way to discriminate the function of TFH memory cells will be to understand where these cells are localized. During the effector phase, TFH cells localize to B cell follicles by interacting with ICOSL expressed by bystander B cells (Shi et al., 2018). If TFH memory cells support the conversion of memory B cells into plasma cells, we would expect this co-localization to continue at late time points after infection. On the other hand, if TFH memory cells directly provide survival signals to plasma cells, they may be localized in or near to the red pulp. In this scenario, TFH memory cell interactions with CD11c+ ICOSL-expressing antigen presenting cells would be impaired by ICOSL blockade, leading to a loss of TFH memory cells (Teichmann et al., 2015).

In summary, our data highlight long-lived TFH memory cells as an attractive target for vaccination. When identified by the absence of CCR7 and CD62L, TFH memory cells can be considered a TEM subset. Indeed, CD4 TEM cells were recently shown to be generated in response to a universal influenza vaccination and correlate with improved cellular and humoral secondary responses following challenge (Valkenburg et al., 2018). In contrast, the unanticipated scarcity of TCM that we observed at very late time points after infection, combined with their more limited recall potential, indicates that vaccines targeting TCM may not offer optimal long-term protection.

## EXPERIMENTAL PROCEDURES

### Mice

Mice were bred and housed under specific pathogen-free conditions at the University Hospital of Basel according to the animal protection law in Switzerland. For all experiments, male or female sex-matched mice were used that were at least 6 weeks old at the time point of infection. The following mouse strains were used: C57BL/6 CD45.2, C57BL/6 CD45.1, Nur77 GFP, T-bet ZsGreen (provided by Jinfang Zhu, NIH, USA), P2X7R^−/−^ (provided by Fabio Grassi, IRB, Switzerland), Bcl6 RFP (provided by Chen Dong, Tsinghua University, China), SMARTA, NIP (Shane Crotty, LJI, USA).

### Infection

Mice were infected by intraperitoneal (i.p.) injection with 2×10^5^ focus forming units (FFU) of LCMV Armstrong or intravenously (i.v.) with 1×10^7^ colony forming units (CFU) ActA deficient *Listeria monocytogenes* (Lm.ActA). In adoptive transfer studies, recipient mice were infected with LCMV Armstrong by i.p. injection with 2×10^5^ on the day following cell transfer. LCMV was grown on BHK-21 cells and titrated on 3T3-cells as described previously (Battegay, 1993).

### In vivo treatments

Anti-ICOSL (HK5.3, Bioxcell, #BE0028) treatment was performed by i.p. injection. Each mouse received 4 injections of 100μg every 72 hours. 2-Deoxy-D-glucose (Sigma, #D8375) was supplied in the drinking water for 14 days at a concentration of 6mg/ml. Rapamycin or vehicle control (1% DMSO in PBS) was given daily by i.p. injection of 75μg/kg body weight for 14 days. For BrdU labelling experiment, BrdU was provided in the drinking water at 0.8mg/ml for 12 days.

### NICD-protector

Mice were intravenously (i.v.) injected with 25μg of commercial (BioLegend, #149802) or 12.5μg homemade ARTC2.2-blocking nanobody s+16 (NICD-protector) 15 minutes prior to organ harvest.

### Mixed bone marrow chimeras and adoptive transfers

Wild-type (WT) host CD45.1 mice were lethally irradiated with 2 fractionated doses of 500 cGγ and reconstituted with a 1:1 mixture of bone marrow cells from CD45.1 WT and P2X7R^−/−^ CD45.2 donor mice. The reconstitution ability of T- and B cells was assessed at least 6 weeks after reconstitution and before LCMV infection. Adoptive transfer experiments with CD4 Smarta and NIP cells were performed as previously described (Moon et al., 2009).

### Quantitative PCR

RNA from sorted cells was isolated with RNAqueous™-Micro Total RNA Isolation Kit (Invitrogen™) and was reverse transcribed to cDNA using either qScript XLT cDNA SuperMix (Quantabio) or iScript™cDNA Synthesis Kit (Bio-Rad). Samples were run on an Applied Biosystems Viia7 Real-time PCR system. Primers used for the indicated genes are listed in the supplementary table.

### Isolation of lymphocytes

Single-cell suspensions of cells were prepared from spleens, blood, liver, and bone marrow. Spleens were either mashed and filtered through a 100μm strainer or digested at 37°C for 1 hour using Collagenase D and DNase I. To isolate bone marrow, femur and tibia bones were flushed with a 25G needle followed by filtration through a 100μm strainer. Livers were perfused with PBS and mashed through a 100μm strainer, followed by gradient centrifugation in Percoll. Lymphocytes from the blood were isolated by gradient centrifugation in Ficoll (LSM MP Biomedicals). Erythrocytes were lysed using Ammonium-Chloride-Potassium (ACK) lysis buffer.

### Flow cytometry and cell sorting

Isolation of LCMV-specific CD4 T cells was performed by staining single-cell suspensions for 1 hour at room temperature with Ia^b^:NP_309-328_ PE or IA^b^:GP_66-77_ APC (provided by NIH tetramer core), followed by enrichment and counting (Moon et al., 2009). Anti-CXCR5 BV421 was added at the time of tetramer staining; for p-S6 detection, cells were stained with tetramer for 20 minutes on ice in the presence of 50nM Dasatinib. LCMV-specific B cells were detected as previously described, using a tetramer for NPΔ340 and a Decoy reagent to discriminate PE-specific B cells (Taylor et al., 2012). All other surface staining was performed for 30min on ice in presence of a viability dye. Transcription factor staining was performed by fixation and permeabilization using the eBioscience Foxp3/Transcription Factor staining set. To assess phospho-protein levels, cells were fixed with BD Phosflow Lyse/Fix Buffer for 10 min at 37°C and permeabilized using BD Phosflow Perm/Wash Buffer for 30 min at room temperature. To stain for BrdU incorporation, the FITC BrdU Flow kit (BD) was used. 2-NBDG staining (Thermo Fisher) was performed for 10 min at 37°C in 100μM 2-NBDG. Fortessa LSR II and Sorp Aria cytometers (BD Biosciences) were used for flow cytometry and cell sorting respectively. Data were analyzed with FlowJo X software (TreeStar).

The following antibodies were used for staining: CD4 (BUV496, GK1.5, BD, #564667), CD8a (biotin, 53-6.7, BioLegend, #100704), CD11b (PE-Cy5, M1/70, BioLegend, #101222), CD11c (PE-Cy5, N418, BioLegend, #117316), CD27 (BV510, LG.3A10, BioLegend, #124229), CD38 (BV421, 90, BD, #562768), CD44 (BUV395, IM7, BD, #740215; APC-Cy7, IM7, BioLegend, #103028; PE, IM7, BD, #553134), CD45.1 (PE, A20, BioLegend, #110707; FITC, A20, BD, #553775), CD45.2 (FITC, 104, BD, #553772; APC-eFluor780, 104, eBioscience, #47-0454-82; APC-Fire, 104, BioLegend, #109852), CD62L (APC, MEL-14, BD, #553152; BV711, MEL-14, BioLegend, #104445), CD69 (FITC, H1.2F3, BD, #553236; PE, H1.2F3, eBioscience, #12-0691-83; APC-Cy7, H1.2F3, BioLegend, #104526; BV785, H1.2F3, BioLegend, #104543), CD73 (APC Fire, TY/11.8, BioLegend, #127221), CD80 (PE-Cy7, 16-10A1, BioLegend, #104734), CD138(BV421, 281-2, BioLegend, #142507; BV711, 281-2, BioLegend, #142519), FAS (PE, 15A7, eBioscience, #12-0951-83; FITC, 15A7, BD, #554257), PSGL-1 (BV605, 2PH1, BD, #740384, BUV395, 2PH1, BD, #740273), CXCR4 (BV711, L276F12, BioLegend, #146517), CCR7 (PE, 4B12, BioLegend, #120106), ICOSL (PE, HK5.3, eBioscience, #12-5985-81), ICOS (PE, 7E.17G9, BioLegend, #117406), PD1 (BV785, 29F.1A12, BioLegend, #135225), IgD (BV510, 11-26c.2a, BioLegend, #405723), IgM (BV786, R6-60.2, BD, #564028), Vα2 TCR (FITC, B20.1, eBioscience, #11-5812-82, PE, B20.1, eBioscience, #12-5812-82), Vβ8.3 (FITC, 1B3.3, BD, #553663; PE, 1B3.3, BD, #553664), B220 (PE-Cy5, RA3-6B2, BioLegend, #103210; BV650, RA3-6B2, BioLegend, #103241; PE-Cy7, RA3-6B2, BioLegend, #103222; biotin, RA3-6B2, BD, #553085), CXCR5 (BV421, L138D7, BioLegend, #145512), F4/80 (PE-Cy5, BM8, BioLegend, #123112), FR4 (PE-Cy7, 12A5, BioLegend, #125012, APC-Fire, 12A5, BioLegend, #125013), GL7 (Alexa Fluor 647, GL7, BioLegend, #144606; Alexa Fluor 48, GL7, BioLegend, #144612), GR-1 (PE-Cy5, RB6-8C5, BioLegend, #108410), Ly6C (BV711, HK1.4, BioLegend, #128037; BV510, HK1.4, BioLegend, #128033; FITC, AL-21, BD, #553104), I-Ab (biotin, AF6-120.1, BioLegend, #116404), NK1.1 (biotin, PK136, BioLegend, #553163), P2X7R (PE, 1F11, BioLegend, #148704; PE-Cy7, 1F11, BioLegend, #148707), Bcl6 (PE-Cy7, K112-91, BD, #563582; PE, K112-91, BD, #561522; AF488, K112-91, BD, #561524), FoxP3 (Alexa Fluor 488, 150D, BioLegend, #320012; PE-Cy7, FJK-16s, eBioscience, #25-5773-82), HIF-1a (PE, 241812, R&D Systems, #IC1935P; Alexa FLuor 700, 241812, R&D Systems, #IC1935N), phospho-AKT S473 (Alexa Fluor 647, D9E, Cell Signalling Technology, #4075S), phospho-S6 S240/244 (Rabbit mAb, Cell Signalling Technology, #5364S), Goat anti-Rabbit IgG (Alexa Fluor 488, Life technologies, #A11034), TCF1 (PE, S33-966. BD, #564217), Zombie Fixable Viability Dye (Zombie Red, BioLegend, #423110; Zombie Green, BioLegend, #423112).

### Enzyme-linked immunosorbent assay

LCMV-specific NP serum antibody titers were determined as previously described (Sommerstein et al., 2015). Briefly, 96-well plates were coated with 100ul of recombinant NP at 3μg/ml in sodium carbonate buffer adjusted to pH 9.6. Plates were blocked with 5% milk in PBS-Tween 0.05% before adding the prediluted sera to the plate. ABTS color reaction was used to detect HRP-coupled goat anti-mouse IgG antibody (Jackson, 115-035-062). Optical density was measured with a Synergy H1 reader (BioTek).

### Detection of LCMV NP-specific IgG antibody secreting cells by ELISpot

LCMV NP-specific antibody secreting cells (ASC) were detected as previously described (Schweier et al., 2019). Briefly, 96-well plates were coated with 100ul of recombinant NP at 3μg/ml in PBS. Plates were blocked with cell culture medium before adding pre-diluted single-cell suspension from the bone marrow or the spleen. Cells were incubated for 5h at 37°C before HRP-coupled goat Fcy-specific anti-mouse IgG antibody was added (Jackson, 115-035-008). AEC solution (BD Bioscience) was used to detect ASC. Spots were quantified using an ELISpot reader from AID.

### scRNA-seq

#### Sample preparation

3-10 × 10^3^ LCMV-specific CD4 memory cells (CD4^+^dump^−^ CD44^+^GP66^+^) from either T-bet ZsGreen (1x sample) or wild-type mice (2x biological replicates) day 35 and day 37 post infection were provided for library preparation using the 10x Chromium platform. Each sample represented cells from 1-3 mice. Single-cell capture and cDNA and library preparation were performed with a Single Cell 3’ v2 Reagent Kit (10× Genomics) according to manufacturer’s instructions. Sequencing was performed on one flow-cell of an Illumina NexSeq 500 at the Genomics Facility Basel of the ETH Zurich. Paired-end reads were obtained and their quality was assessed with the FastQC tool (version 0.11.5). The length of the first read was 26 nt, composed of individual cells barcodes of 16 nt, and unique molecular identifiers (UMIs) of 10 nt. The length of the second read, composed of the transcript sequence, was 58 nt. The samples in the different wells were identified using sample barcodes of 8 nt.

#### Sample analysis

Sequencing data was processed using 10X Genomics’ cellranger software version 2.1.0, modified to report only one alignment (randomly) for multi-mapped reads (keep only those mapping up to 10 genomic locations). Raw molecule info from cellranger was initially filtered leniently, discarding cells with fewer than 100 genes. The resulting UMI matrix was further filtered to keep only cells with log library size > 2.8, log number of features > 2.6, % mitochondrial reads <= 6, % ribosomal protein reads >= 20. Genes with average counts < .005 were removed. Normalization was done using the R package scran’s deconvolution method, where cells are pre-clustered, normalized within each cluster, and normalized between each cluster. Technical noise within gene expression was modeled using scran, and biologically relevant highly variable genes were calculated after separating the technical from biological variance, using FDR < 0.05 and biological variance > 0.1. A PCA was run on the normalized data using the top 500 most variable genes by biological variance, and the PCA was denoised to account for the modelled technical variation. Cells were clustered hierarchically using Ward’s method on the distance matrix of the PCA. Default dendrogram cut height using the R package dynamicTreeCut resulted in 3 clusters, which also mapped to the highest average silhouette width. The cut height for a finer clustering (7) was chosen based on the next highest local maximum in a plot of average silhouette width. *t-*distributed stochastic neighbor embedding (tSNE) was executed with a perplexity of 30. GSEA between clusters was performed using camera from the limma R package on standard gene set categories (MSigDB) as well as sets curated from relevant publications (Liberzon et al., 2011; Ritchie et al., 2015; Subramanian et al., 2005). Subsequent visualization and analysis such as dropout imputation were performed using version 2.3.4 of the Seurat R package. Single-cell regulatory network interference and clustering (SCENIC) was performed using version 0.9.7 and the published workflow on the pySCENIC github repository (Aibar et al., 2017). Pseudotime analysis was performed with Monocle2 version 2.10.1. Normalized data as above were further filtered to exclude genes found in less than 5% of cells. Dimension reduction was performed using DDRTree and differentially expressed genes calculated with the differentialGeneTest function based on the hierarchical clustering above. The root state of the trajectory was chosen where Tcf7 expression was highest.

In silico pseudo-bulk data was generated from three biological replicates (one without and two with NICD-protector). Biological replicate, percent mitochondrial reads and percent ribosomal reads each explained < 0.1 % of variance in the data. Counts for each gene were summed by cluster, and data re-normalized using Trimmed Mean of M-Values (TMM). Differential expression analysis was performed between the most extreme TFH and Th1 clusters using edgeR and limma. Ingenuity Pathway Analysis was performed on the resulting list of differentially expressed genes (adj P-value < 0.05).

### ATAC-seq

#### Sample preparation

6.5-7.5 × 10^3^ LCMV-specific TFH cells (Ly6C^lo^PSGL1^lo^), TH1 cells (Ly6C^hi^PSGL1^hi^) were sorted in duplicates from different mice to obtain biologically independent samples 32 days post infection. Naïve CD4 cells (CD44^lo^ CD62L^hi^) were sorted in duplicates and represent technical replicates. Library preparation for sequencing was performed as described previously with an additional clean up step to reduce reads mapping to mitochondrial DNA(Corces et al., 2017). Briefly, sort-purified cells were lysed and tagmented for 30 min at 37°C. DNA was purified using Zymo DNA Clean and Concentrator Kit and amplified for five cycles. Real-time PCR was used to determine the number of additional PCR cycles. DNA was cleaned up using AMpure XP beads at a 1.2 × ratio twice. Sequencing was performed on an Illumina NexSeq 50 machine using 41-bp paired-end run.

#### Alignment and basic QC

Reads were aligned to the mouse genome (UCSC version mm10) with bowtie2 (version 2.3.2) using options “--maxins 2000 --no-mixed-- no-discordant --local --mm”. The output was sorted and indexed with samtools (version 1.7) and duplicated reads were marked with picard (version 2.9.2). The read and alignment quality was evaluated using the qQCReport function of the Bioconductor package QuasR (R version 3.5.2; Bioconductor version 3.8).

#### Single sample peak calling

For each group of biological replicates, regions of accessible chromation where called with macs2 (version 2.1.1.20160309) using the option ‘-f BAM -g 2652783500 --nomodel --shift -100 --extsize 200 --broad --keep-dup all --qvalue 0.05’. Since the majority of duplicated reads came from mitochondria, the resulting peak lists were cleaned from peaks called in mitochondria and additionally ENCODE blacklist regions. All peaks also required a log-fold change > 1 and FDR < 0.05. The resulting peak lists, one per population, were merge using the function reduce from the GenomicRanges package, requiring at least 250 bases gap for separate peak calls and a minimum peak width of 100 bases. This merged peak list resulted in 118999 peaks (5.1% genome) across all five populations. The Bioconductor package bamsignals was used to generate a counts matrix for the merged peak list.

#### Across sample normalization and peak calling

TFH and Th1 samples differed strongly from naïve samples in the enrichment of reads within accessible chromatin as compared to genomic background (Supplementary Methods Figure A). One possible source of this bias is a differential tagmentation efficiency between samples or cell types as previously observed (Denny et al., 2016). This bias also leads to an artificial regulation when comparing ATAC signals of open versus closed promoters (Supplementary Methods Figure B). Here we do not expect a difference in the dynamic range between both states across samples/cell types. In order to compare log-fold changes across samples we therefore applied quantile normalization to log-CPM levels of the TMM-scaled peak counts. The resulting peak intensities were converted to a Bioconductor ExpressionSet object and all further differential accessibility analysis was performed with this object. Specifically, the limma package was used for differential accessibility between all pairs of populations using functions lmFit and eBayes

#### Differential motif analysis using HOMER

Fasta sequences of Th1 and TFH peaks were extracted using functions from the BSgenome, Biostrings and rtracklayer R/Bioconductor packages. These sequences were stratified according to population specific peaks (=open in only one population) and common peaks (= open in both) based on thresholding the differential accessibility results by logFC > 2 and FDR < 0.05. Homer (version 4.9) was used to predict known TF motifs (motif database: vertebrates/known.motif) within these sequence sets by running the command ‘homer2 known-strand both’ with additional setting specifying the specific peaks as foreground and the common peaks as background sequence sets. Differential motif occurrence results were visualized using R.

#### Ingenuity pathway analysis

Ingenuity pathway analysis (Qiagen, version 01-14) was performed on differentially accessible peak regions (log_2_ FC > 0.5 and FDR of 0.05) in Th1 vs TFH memory cells with default settings.

### Bulk RNA-seq

#### Sample preparation

0.5-4.0 × 10^3^ GP66-specific TFH cells (Ly6C^lo^PSGL1^lo^) were sorted in quadruplicates from individual mice to obtain biologically independent samples from control and αICOSL treated mice. Total RNA was isolated with the PicoPure™ RNA Isolation kit (ThermoFisher, # KIT0202). cDNA and library preparation were performed with a SMART-Seq v4 Ultra Low Input RNA Sequencing Kit (Takara Bio) according to manufacturer’s instructions. Sequencing was performed on one flow-cell of an Illumina NexSeq 500 using 38-bp paired-end run.

#### Sample analysis

Single-end RNA-seq reads were mapped to the mouse genome assembly, version mm10 (analysis set, downloaded from UCSC https://genome.ucsc.edu), with STAR (version 2.7.0c, (Dobin et al., 2013)), with default parameters except for reporting only one hit for multi-mappers in the final alignment files (outSAMmultNmax=1) and filtering reads without evidence in spliced junction table (outFilterType=“BySJout”).“) (Dobin et al., 2013). All subsequent gene expression data analyses were done within the R software (R Foundation for Statistical Computing, Vienna, Austria). Using Ensembl Genes mRNA coordinates from ensembl version 84 (https://www.ensembl.org) and the featureCounts function from Rsubread package, we quantified gene expression as the number of reads that started within any annotated exon of a gene (Liao et al., 2019). Only genes annotated as “protein_coding” were kept for further analysis. Genes that did not show an expression level of at least 0.5 CPM in each of the 8 samples were filtered out and TMM normalization performed. Differential expression analysis was performed with the Bioconductor limma package using the function voomWithQualityWeights and applying a cyclic-loess normalization. We found this methodology to perform best in dealing with the reduced transcriptome complexity in samples with the lowest number of cells used as input. Although no gene was significantly differentially expressed at an FDR threshold of 5%, the top genes were consistent with protein expression experiments. Gene set enrichment analysis was performed using the cameraPR function from limma on standard gene set categories (MSigDB) as well as sets curated from relevant publications.

### Extracellular metabolic flux analysis

A Seahorse XFe96 metabolic extracellular flux analyzer was used to determine the extracellular acidification rate (ECAR) in mpH/min and the oxygen consumption rate (OCR) in pmol/min. In brief, T-cells were sorted in CD4+CD44+ TFH cells (Ly6C^lo^PSGL1^lo^) or TH1 cells (Ly6C^hi^PSGL1^hi^) and seeded (2×10^5^/well at memory time point or 2.5 x10^5^/well at effector time point in respective experiments) in serum-free unbuffered RPMI 1640 medium (Sigma-Aldrich # R6504) onto Cell-Tak (#354240, Coring, NY, USA) coated cell plates. Mito Stress test was performed by sequential addition of oligomycin (1 μM; Sigma Aldrich 75351), carbonyl cyanide-4-(trifluoromethoxy)phenylhydrazone (FCCP; 2 μM; Sigma Aldrich C2920) and rotenone (1 μM; Sigma Aldrich R8875) at the indicated time points. Metabolic parameters were calculated as described previously (Gubser et al., 2013).

### Statistical analysis

For statistical analysis of one parameter between two groups, unpaired two-tailed Student’s t-tests were used to determine statistical significance. In time course experiments, statistical analysis was performed for each individual timepoint. To compare one parameter between more than two groups, one-way analysis of variance (ANOVA) was used followed by Turkey’s post-test for multiple comparisons. * = *P* <0.05, ** = *P* < 0.01, *** = *P* <0.001, **** = *P* < 0.0001, ns = not significant. Error bars show SD centered on the mean unless otherwise indicated. Data was analysed using GraphPad Prism software (version 7 or 8).

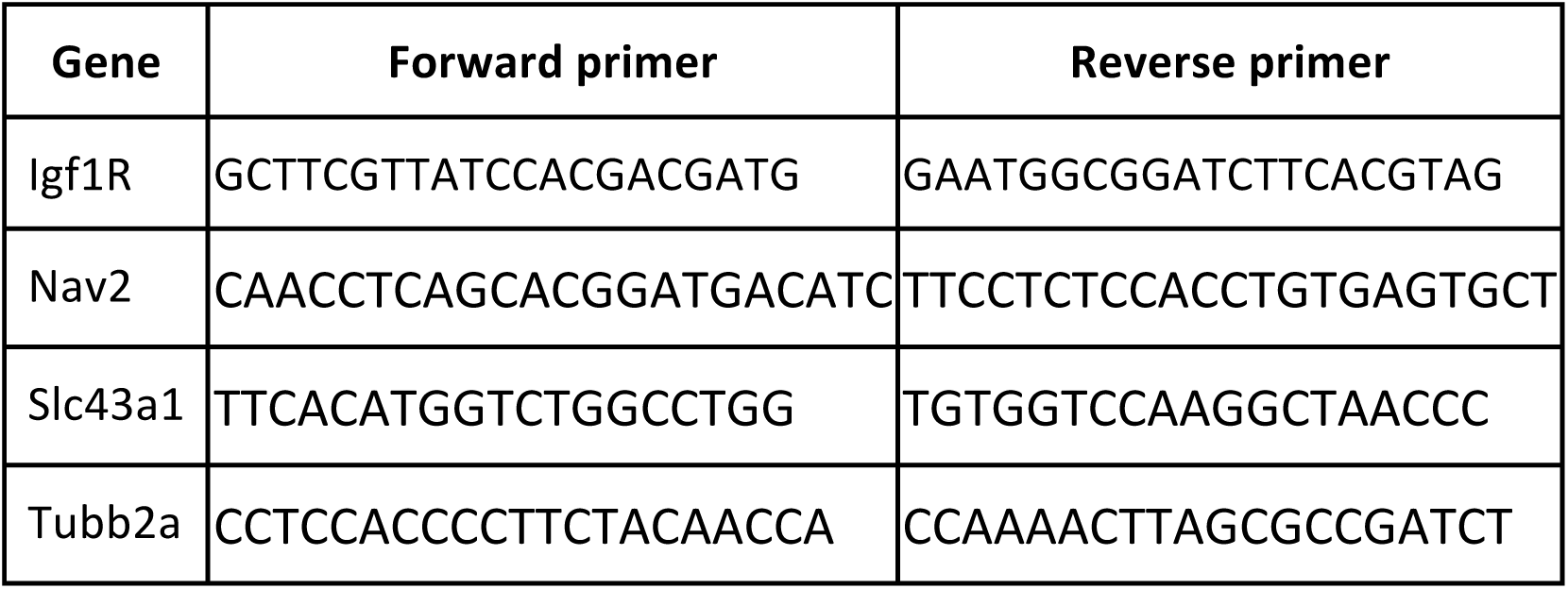
Table for qPCR primers.

## Acknowledgements

We thank D. Pinschewer for many helpful discussions, sharing reagents and technical advice, M. Linterman for critical reading of the manuscript, G. Bantug and D. Labes for expertise and discussion, C. Beisel for technical advice, R. Tussiwand and her lab for feedback and support; R. Jakob, T. Maier, T. Rezzonico and F.Grassi for technical support, the flow sorting facility and all the animal caretakers at the DBM University of Basel. Single-cell RNA-sequencing was performed at the Genomics Facility Basel, ETH Zurich. Calculations were performed at sciCORE (http://scicore.unibas.ch/) scientific computing center at the University of Basel.

## Author contributions

Conceptualization, CK, MK, DS; Methodology, MK, DS, CK; Investigation, MK, TP, CK, JL, NS; Formal analysis, MK, DS, CK, JR, FG; Software, DS, JR, FG; Writing - CK, DS, MK; Visualization, DS, MK, CK; Funding acquisition, CK; Resources, DP, JT, CH; Supervision, CK.

## Data and materials availability

scRNA-seq, ATAC-seq and bulk RNA-seq data are deposited with the National Center for Biotechnology Information Gene Expression Omnibus (accession nos: PLACEHOLDER). NIP-transgenic mice were generated in the lab of Shane Crotty. Requests for this strain should be addressed to Shane Crotty. Bcl6-RFP mice were generated in the lab of Chen Dong. Requests for this strain should be addressed to Chen Dong.

## Supplementary Figure Legends

**Supplementary Figure 1.**
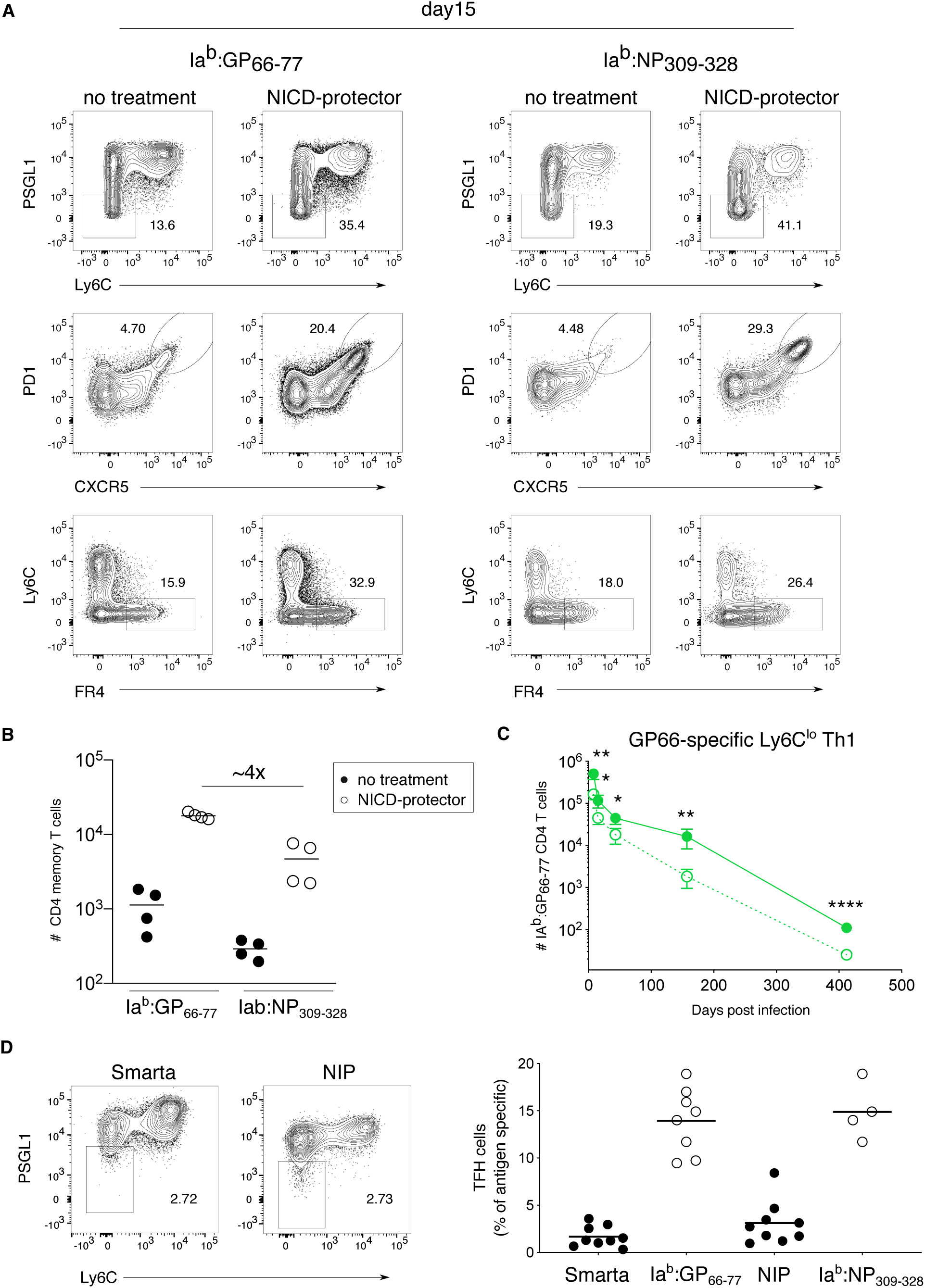
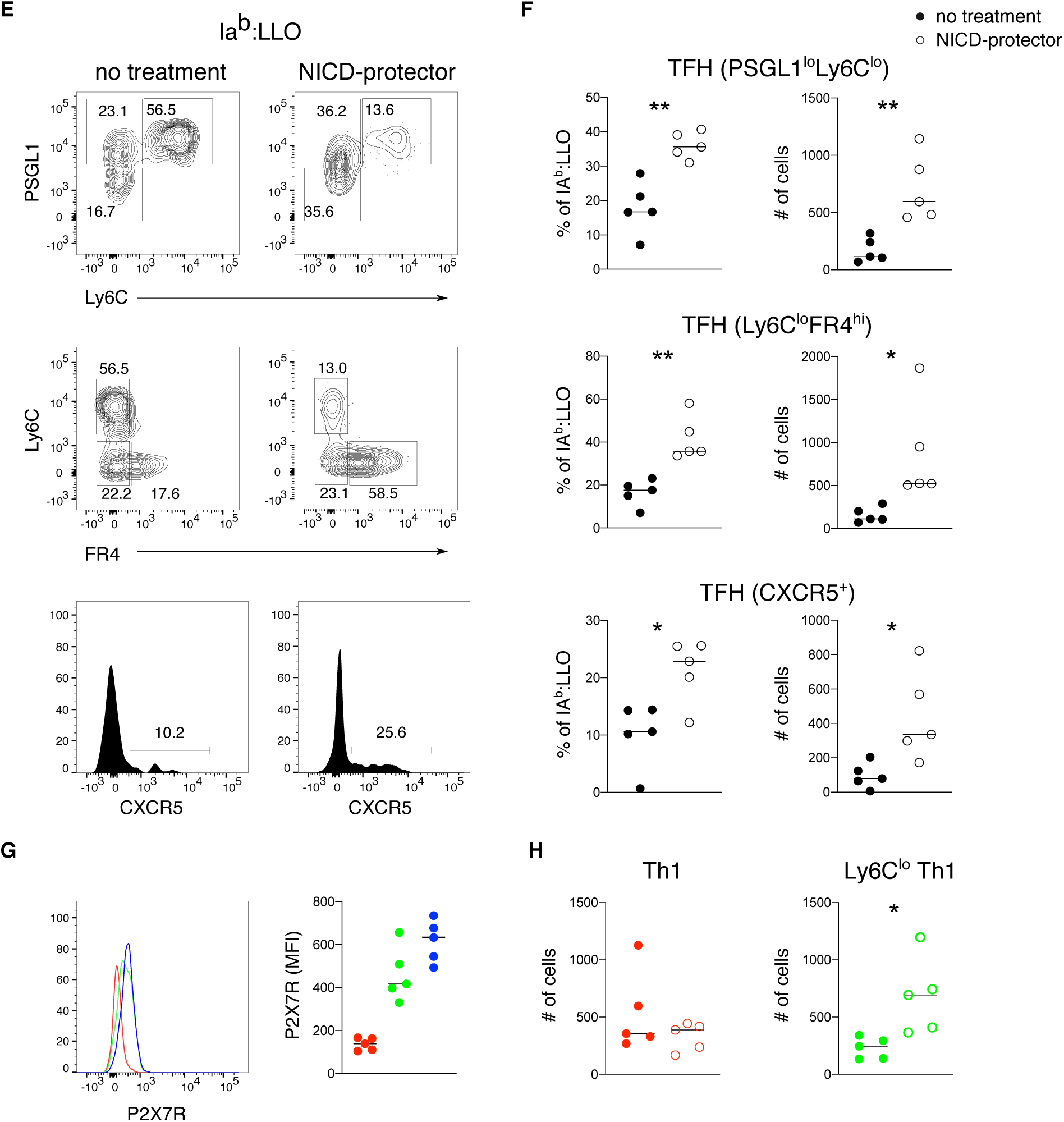
TFH memory cells are susceptible to death during isolation. (A) Flow cytometry plots of splenic CD4 T cells isolated at day 15 post infection, with or without NICD-protector comparing GP66 epitope to NP epitope with different gating strategies to identify TFH memory cells. (B) Comparison of epitope-specific CD4 memory cell numbers in control and NICD-protector treated mice >30 days post infection. (C) GP66-specific Ly6C^lo^ Th1 memory cell numbers (green, gated as in Figure 1D) over time with (solid line) or without (dashed line) NICD-protector. (D) Flow cytometry plots (left) and proportions of TFH memory cells (right) in Smarta and NIP memory cells compared to polyclonal memory compartments >30 days post infection. (E-F) Flow cytometry plots and quantification (F) of splenic LLO-specific TFH cells isolated >30 days post infection with or without NICD-protector after Listeria infection. (G) Histogram (left) and MFI (right) of P2X7R in LLO-specific Th1 (red), Ly6C^lo^ Th1 (green) and TFH (blue) memory cells >30 days post infection. (H) Quantification of LLO-specific CD4 memory subsets isolated >30 days post infection with or without NICD-protector after Listeria infection. Thin lines represent the mean ± s.d. Data represent *(N)* = 2 (B, E, F, G, H) independent experiments with *n* = 3-4 mice or summarize *(N)* = 5 independent experiments (D) with *n* = 4-9 mice per group (dots represent cells from individual mice, and the line represents the mean). Unpaired two-tailed Student’s t test was performed for each individual time point (C). \**P* < 0.05, \*\**P* < 0.01, \*\*\**P* < 0.001.

**Supplementary Figure 2.**
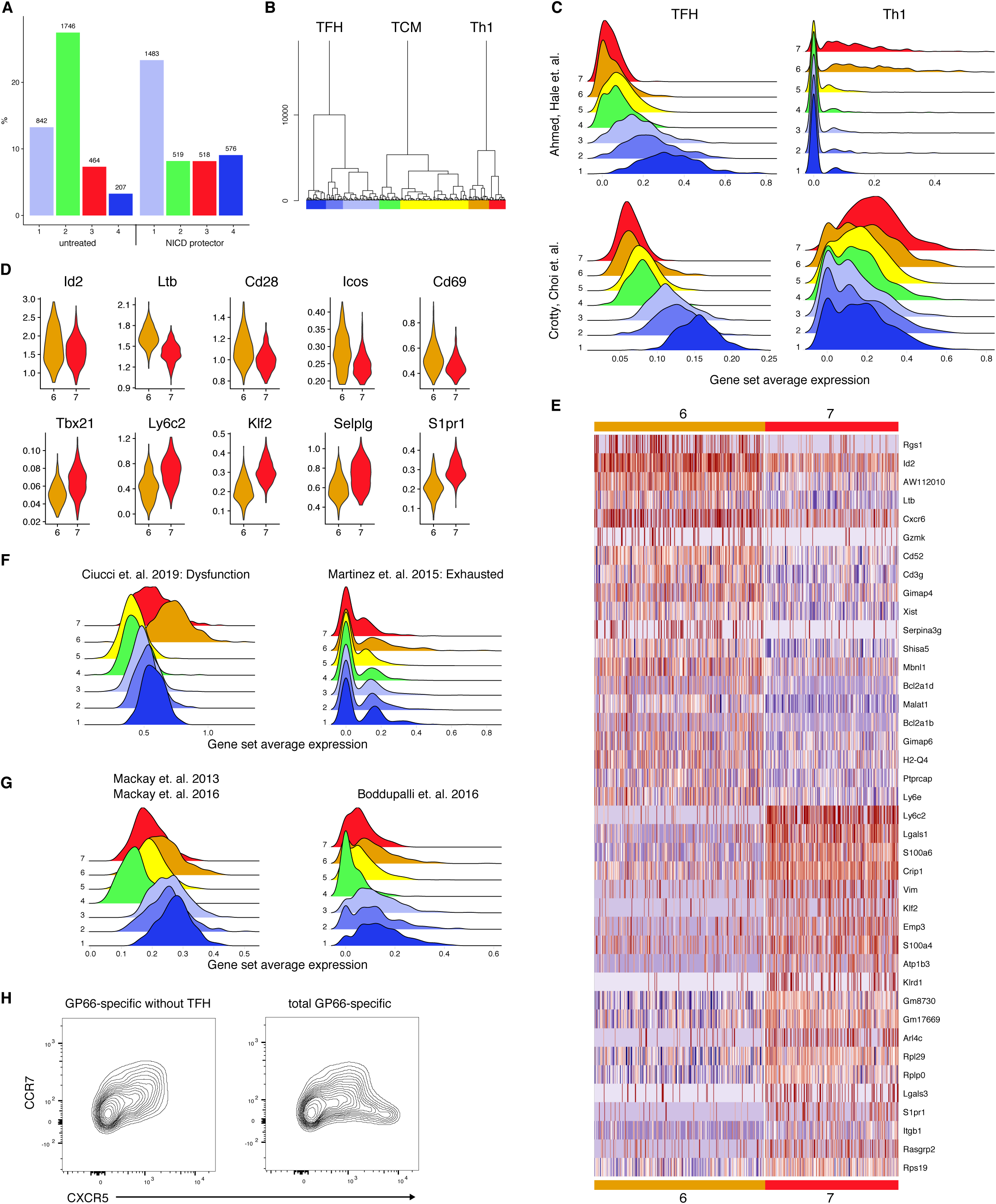
TFH memory cells are transcriptionally distinct from TCM. (A) Clustering analysis of combined scRNA-seq datasets. TFH-like clusters in blue (1, 4), TCM-like in green (2) and Th1-like in red (3). (B) Dendrogram showing hierarchical clustering. Single cells colored by cluster on x axis with tree height on y axis. (C) Log-normalized average expression of published gene sets defining subsets. (D) Selected genes differing between two Th1-like clusters. (E) Heatmap showing top 20 genes segregating Th1-like clusters. (F) Log-normalized average expression of dysfunction (left) (Ciucci et al., 2019) and exhaustion (right) genes (Martinez et al., 2015). (G) Log-normalized average expression of published TRM signatures (Boddupalli et al., 2016; Mackay et al., 2016; Mackay et al., 2013). (H) Flow cytometry analysis of total GP66-specific CD4 memory cells with (right) or without (left) TFH (Ly6CloPSGL1lo). Data represents one of *(N)* = 2 independent experiments with *n* = 4 mice.

**Supplementary Figure 3.**
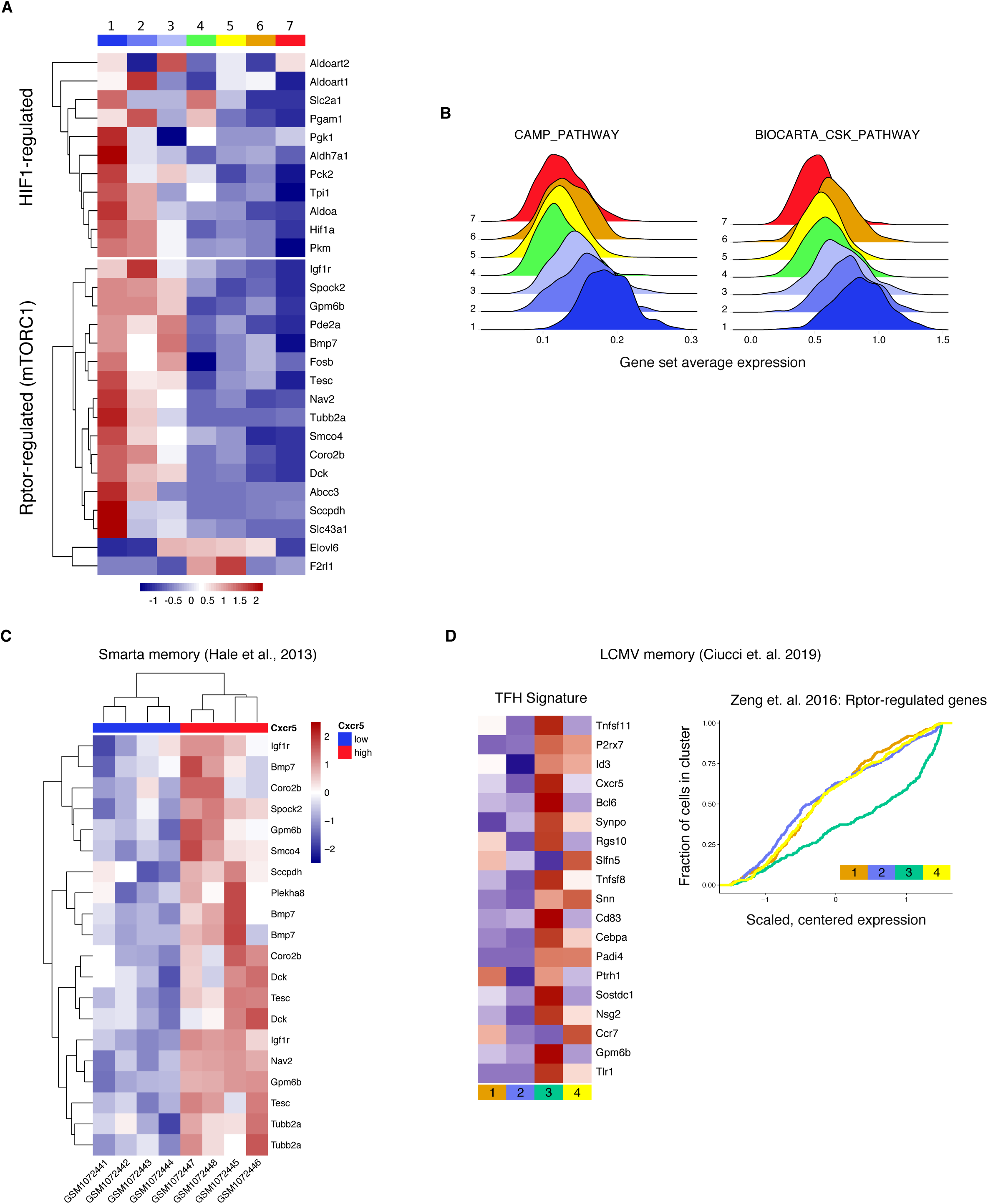
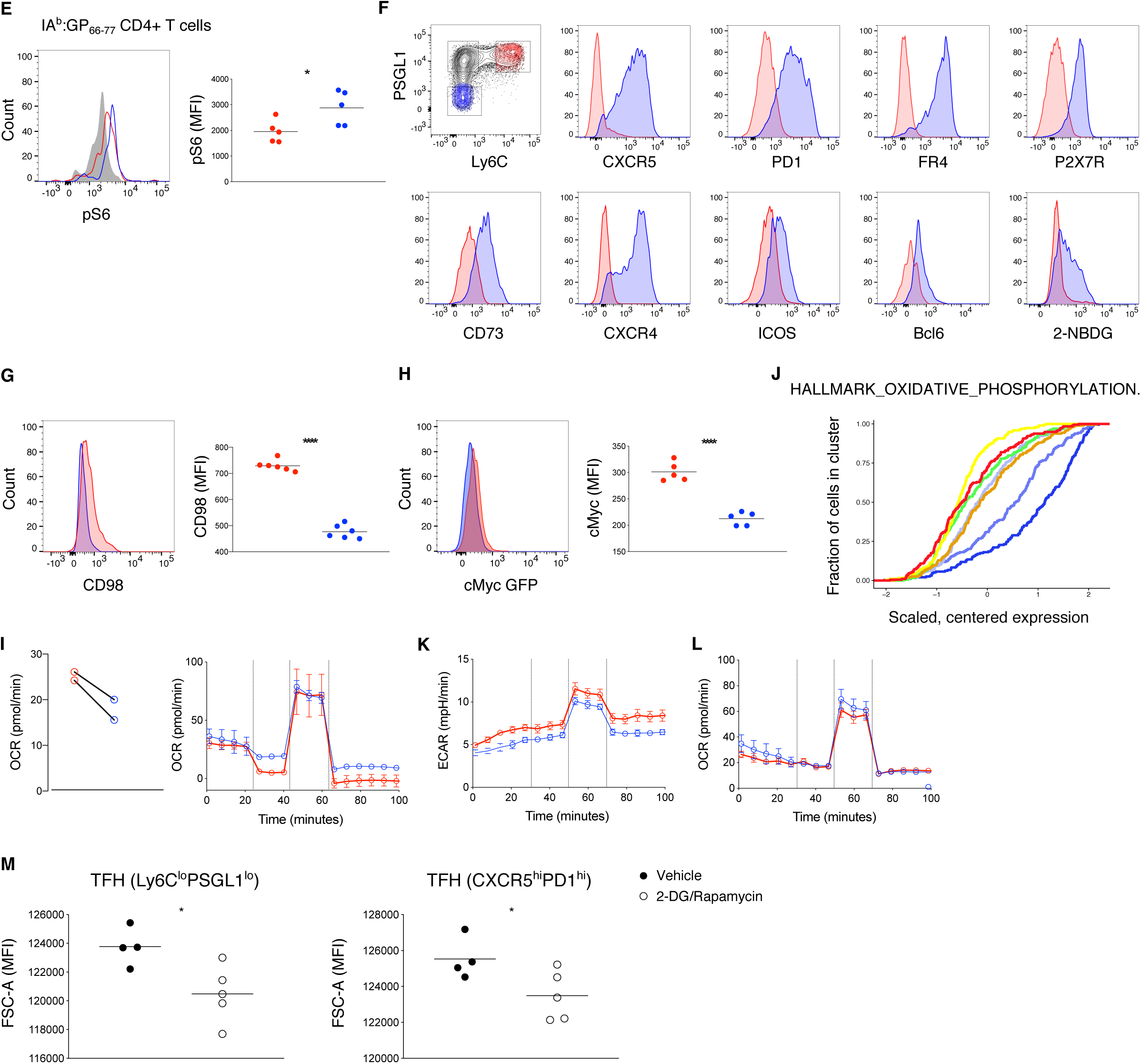
TFH memory cells are constitutively glycolytic. (A) Heatmap showing scaled, centered, per-cluster expression of leading edge mTORC1-related genes (Zeng et al., 2016) and HIF1-regulated genes (Finlay et al., 2012). (B) Log-normalized average expression of CAMP_PATHWAY and BIOCARTA_CSK_PATHWAY gene sets. (C) Analysis of published data from Cxcr5- and Cxcr5+ Smarta CD4 memory cells (Hale et al., 2013) showing leading edge Rptor-regulated genes from Zeng et. al (Zeng et al., 2016). (D) Analysis of published data from d30 CD4 memory to LCMV: heatmap of expression of TFH signature genes and cumulative distribution of Rptor-regulated genes (Ciucci et al., 2019). (E) Flow cytometry analysis of phospho-S6 Ser240/244 and MFI in GP66-specific memory subsets Th1 (red) and TFH (blue) compared to naive CD4 T cells (grey). (F) Flow cytometry analysis of indicated markers in CD44^+^ Th1 and TFH memory cells. (G-H) Histogram and MFI of CD98 (G) or cMyc GFP (H) in GP66-specific memory Th1 and TFH. (I) Quantification of basal respiration and OCR profile in response to Mito-Stress test on sorted CD44^+^ Th1 and TFH memory. (J) Cumulative distribution of gene set HALLMARK_OXIDATIVE_PHOSPHORYLATION. (K-L) ECAR profile (K) and OCR profile (L) in response to Mito-stress test in Th1 and TFH effector cells (day 10 post infection) pooled from 12-14 mice. (M) MFI of FSC-A in GP66-specific TFH memory cells treated with vehicle or 2-DG/rapamycin. *(N)* = 2 independent experiments with *n* = 5 (E,H,M) or 6 (G) mice. Line depicts the mean and dots cells from individual mice. Unpaired two-tailed Student’s t test was performed with \**P* < 0.05, \*\*\*\**P* < 0.0001.

**Supplementary Figure 4.**
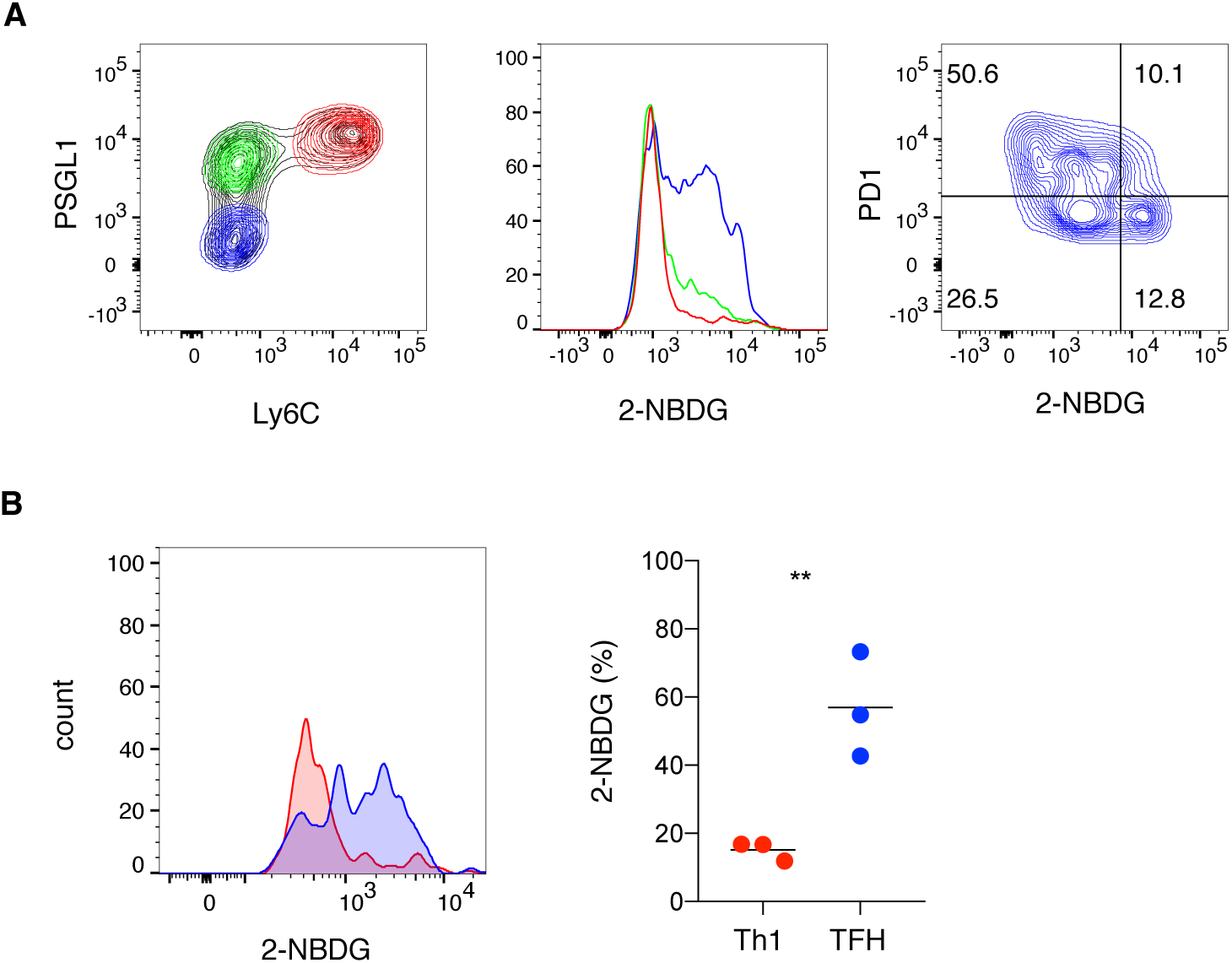
TFH cells can survive in the absence of antigen. (A) Representative flow cytometry plot of 2-NBDG uptake (middle) in the different LCMV GP66-specific CD4 memory subsets Th1 (red), Ly6C^lo^ Th1 (green), and TFH (blue). 2-NBDG uptake in the TFH memory compartment (right) versus PD-1 expression. (B) Representative flow cytometry plot of 2-NBDG uptake (left) and proportion of 2-NBDG+ GP66-specific Th1 (red) or TFH (blue) memory cells (right) 412 days post infection. The line represents the mean and each dot represents cells from individual mice. Data are representative of *(N)* = 2 independent experiments with *n* = 3 (B) or 5 (A) mice. Unpaired two-tailed Student’s t test was performed with \*\**P* < 0.01.

**Supplementary Figure 5.**
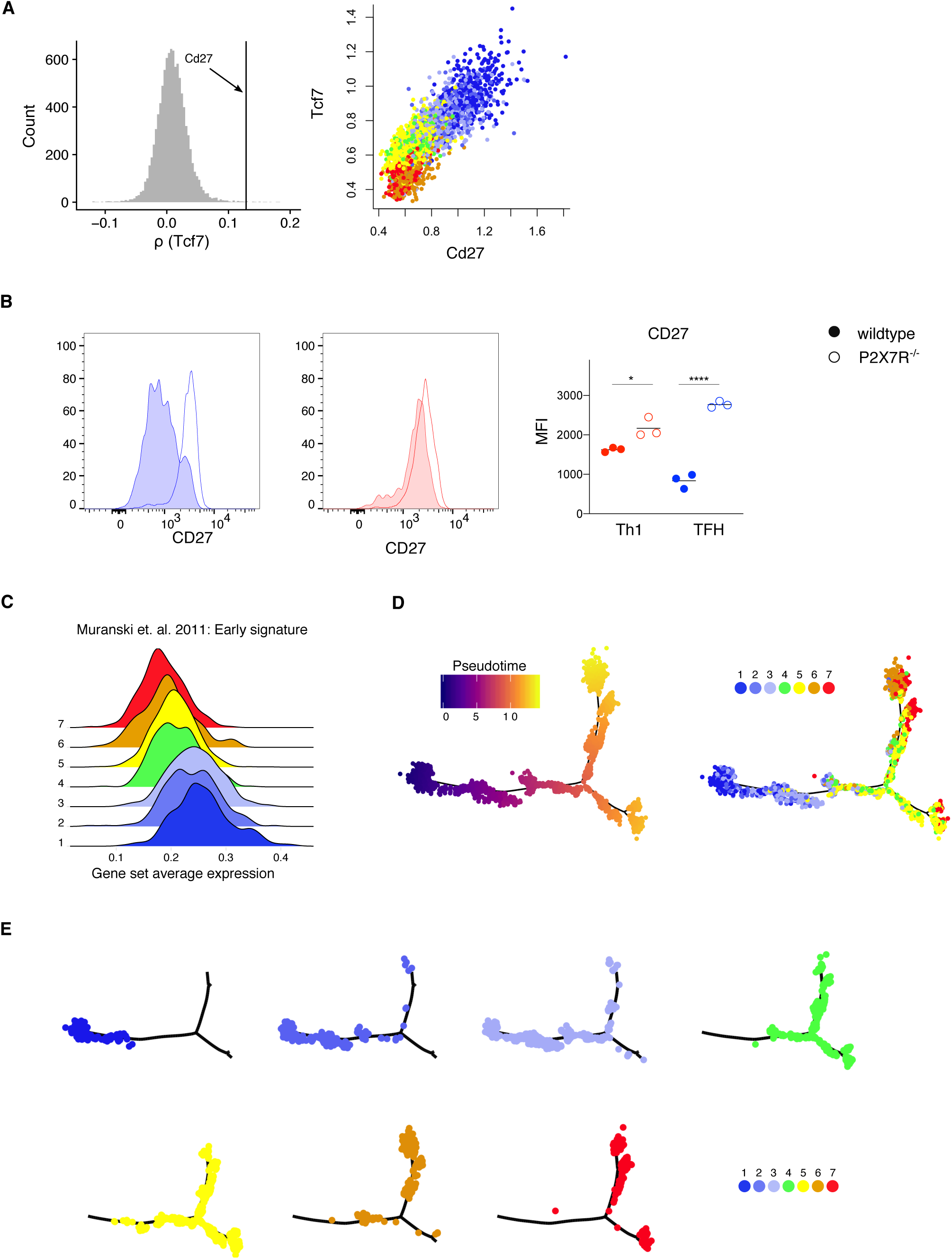
TFH memory cells generate multiple cell fates upon recall. (A) scRNA-seq: Rank of Cd27 among Spearman’s rank correlation coefficients between Tcf7 and all other genes (left). Cd27 vs Tcf7 on imputed data (right), colored by cluster. (B) Flow cytometry analysis (left, middle) and quantification (right) of CD27 expression in P2X7R^−/−^ BM chimera mice in GP66-specific TFH (left) and Th1 memory cells (middle). Filled histograms represent wildtype cells, non-filled histograms depict P2X7R^−/−^ cells. The line depicts the mean and dots represent cells from individual mice. (C) Normalized average scRNA-seq expression of early memory signature (Muranski et al., 2011). (D) Monocle2 analysis showing trajectory in pseudotime (left) and with cluster assignment (right). (E) Monocle2 trajectory with clusters separated. Data represent one of *(N)* = 2 independent experiments with *n* = 3-4 mice (B). Unpaired two-tailed Student’s t test was performed with \**P* < 0.05, \*\*\*\**P* < 0.0001.

**Supplementary Figure 6.**
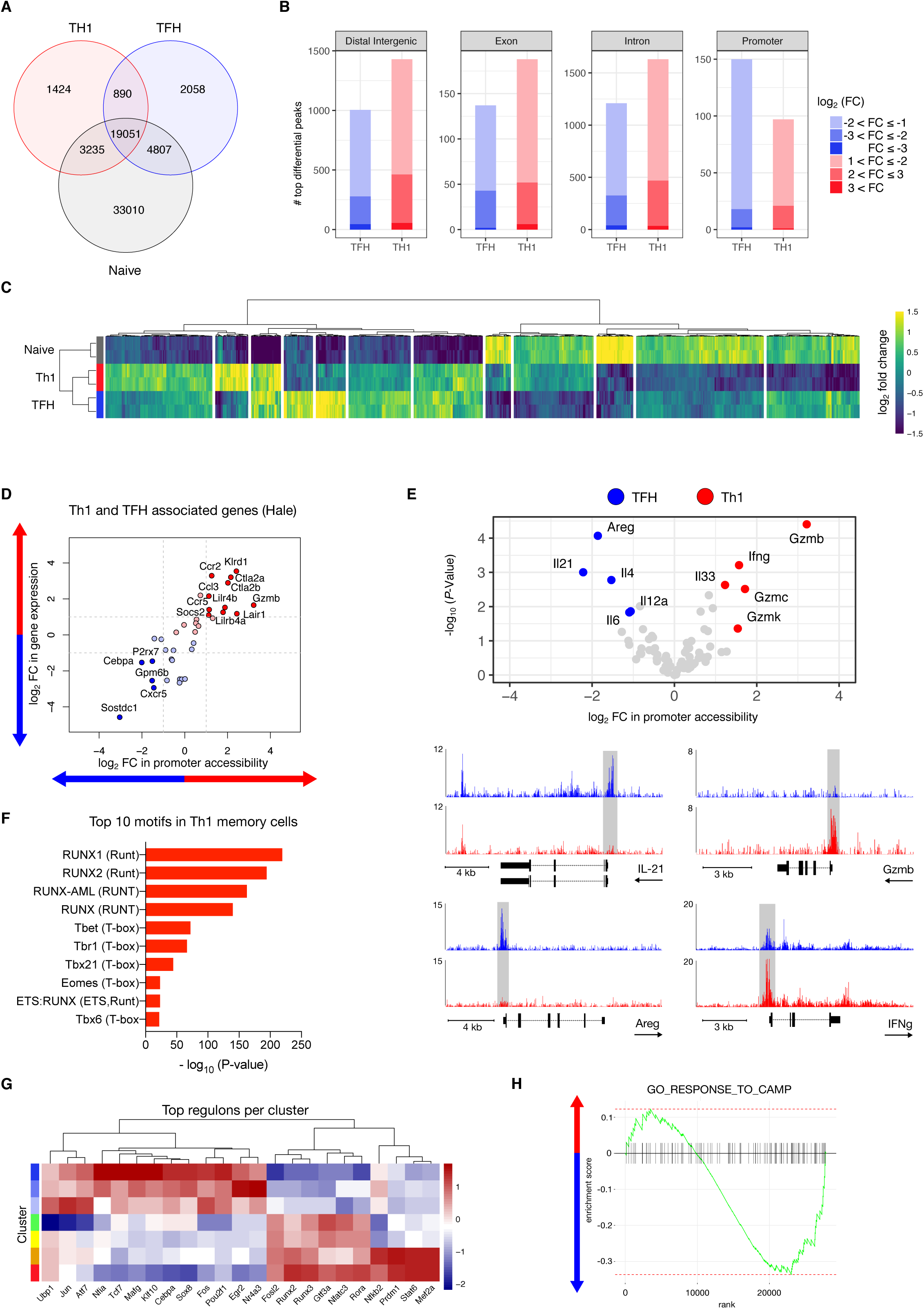
Epigenetic regulation of TFH memory T cells. (A) Venn diagram of all called peaks in Assay for Transposase-Accessible Chromatin using sequencing (ATAC-seq) data depicting the overlap of peaks in the subsets. (B) Number of differentially accessible regions for each genomic feature. (C) Heatmap of hierarchically clustered promoter regions with absolute value (log_2_FC) > 1 in at least one comparison and FDR < 0.1. (D) Scatter plots of ATAC-seq promoter log2 FC and in silico pseudo-bulk RNA-seq log2 FC with blue and red arrows indicating enrichment in TFH and Th1 compartment respectively. (E) Volcano plot of cytokine promoter region accessibility with blue and red dots indicating increased accessiblity in TFH and Th1 compartment respectively and highlighted gene tracks. (F) Enriched motifs in Th1 memory cells found by HOMER. (G) Heatmap depicting top transcriptional network regulators (regulons) on scRNA-seq data found by pySCENIC (Single-Cell rEgulatory Network Inference and Clustering) algorithm. (H) GSEA analysis of the GO_RESPONSE_TO_CAMP pathway with blue and red arrows indicating enrichment in TFH and Th1 respectively.

**Supplementary Figure 7.**
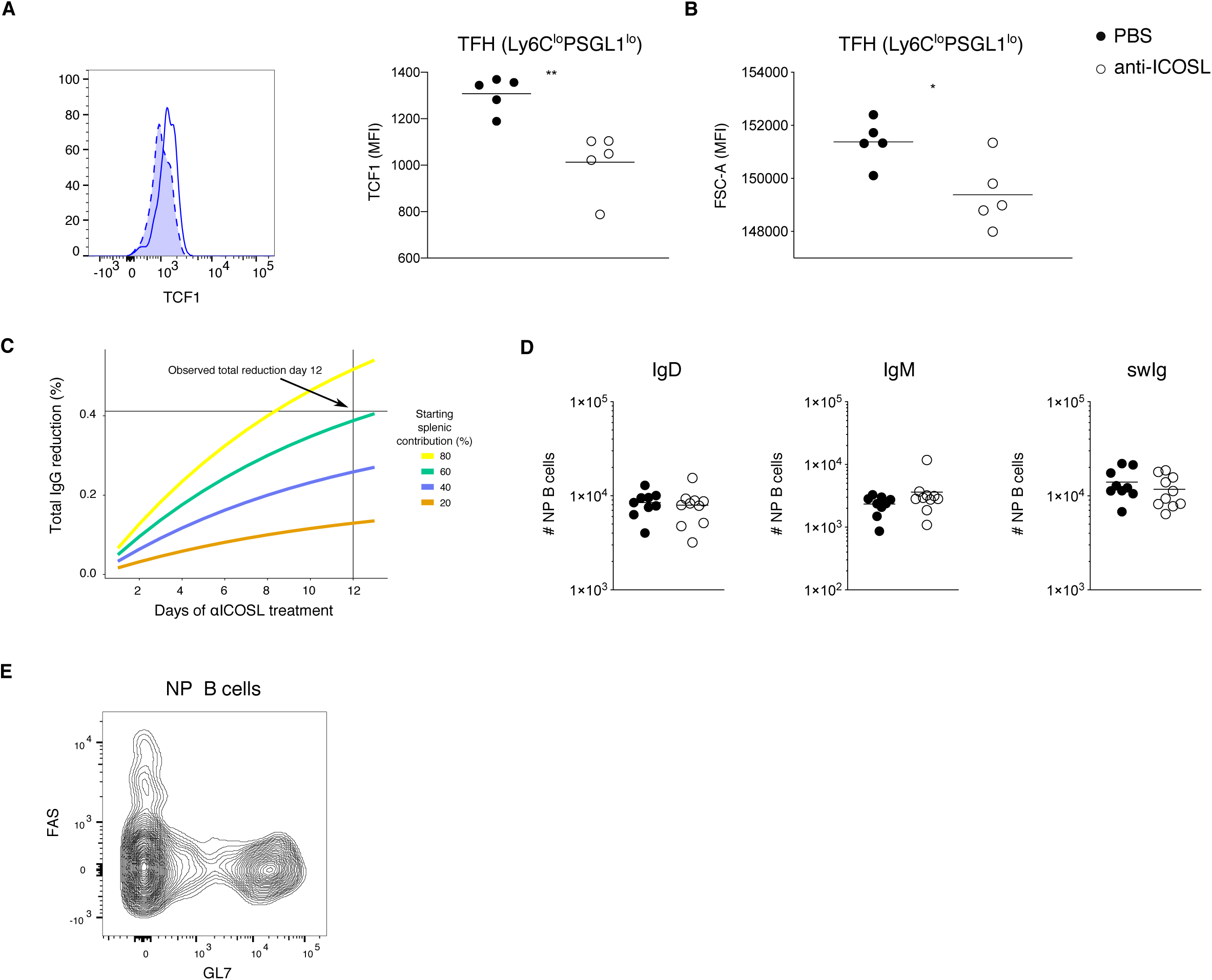
ICOS signaling maintains TFH memory cell identity. (A) Flow cytometry analysis (left) and MFI of TCF1 expression in GP66-specific TFH memory cells comparing PBS (solid line) and anti-ICOSL treated mice (dashed line with filled histogram). (B) MFI of FSC-A in GP66-specific TFH memory cells comparing PBS vs anti-ICOSL treated mice. The thin line represents the mean and each dot represents cells from an individual mouse. (C) Exponential decay model showing IgG reduction after 12 days of anti-ICOSL treatment at various starting splenic contributions and assuming 100% abrogation of splenic contribution. (D) Cell numbers of IgD, IgM or swIg NP-specific memory B cells comparing PBS and anti-ICOSL treated mice. (E) Flow cytometry analysis of NP-specific B cells. Data is representative of *(N)* = 2 independent experiments with *n* = 5 mice per group (A-B) or summarize *(N)* = 2 independent experiments with *n* = 9-10 mice per group (D-E). The thin line represents the mean and each dot represents cells from an individual mouse. Unpaired two-tailed Student’s t test was performed with \**P* < 0.05, \*\**P* < 0.01.

**Supplementary Methods Figure.**
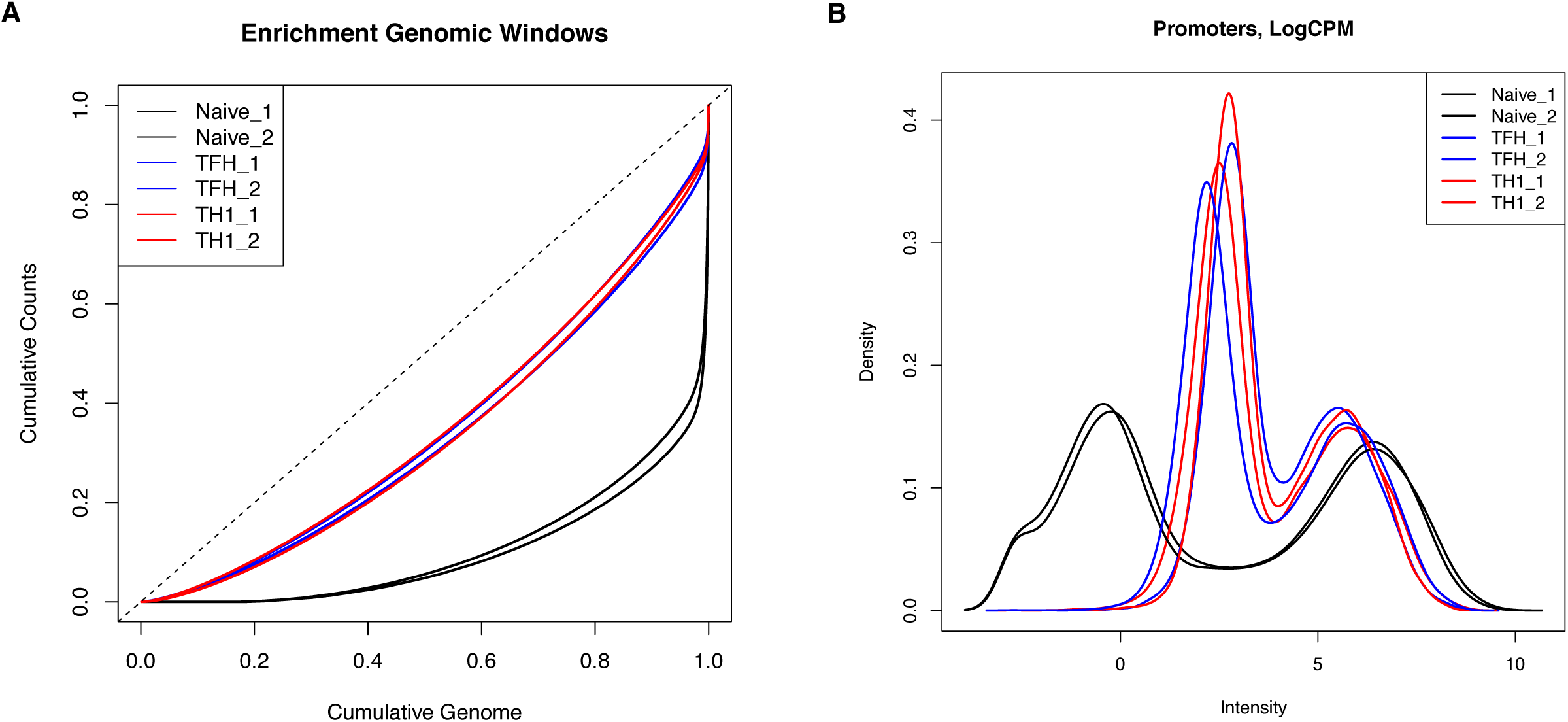
Normalization of ATAC-seq data. (A) Enrichment of genomic windows in the different ATAC-seq samples. (B) Density of logCPM values for the different ATAC-seq samples in promoter regions.

